# Multiplexing of visual-auditory signals in a predominantly auditory brain region

**DOI:** 10.1101/2025.01.28.635385

**Authors:** Meredith N. Schmehl, Yunran Chen, Surya T. Tokdar, Jennifer M. Groh

## Abstract

A recently proposed theory of neural representation postulates that when more than one stimulus is to be encoded, neurons in that representation may fluctuate between encoding each item. To date, such work has considered only cases in which both stimuli are of the same sensory modality, such as how two sounds are encoded in an auditory brain region. Here, we consider the influence of sensory modality on this process, asking whether stimuli of a non-dominant sensory modality can evoke or influence the fluctuating activity patterns observed for combinations of stimuli of a dominant sensory modality. Specifically, we investigated the impact of combining visual and auditory stimuli on single-unit activity in the inferior colliculus, a predominantly auditory structure, in monkeys performing a behavioral task involving reporting all stimulus locations in a given trial. We found that fluctuating activity is evident when a single sound and a single visual stimulus are paired at the same spatial location, indicating that fluctuating activity can be induced when one item is not from a brain area’s preferred modality, and even when only a single spatial location is involved and spatial attention is directed to only one place. Fluctuating activity was also identified when visual stimuli were paired with sounds presented at two spatial locations. The IC appeared to maintain a representation of all stimuli, regardless of the sequence in which the monkey localized the targets (although in one monkey there was a bias toward greater representation of the second reported location). Together, these results indicate that fluctuating activity occurs in response to multimodal stimuli in a predominantly unimodal brain structure and is geared toward preservation of all sensory information rather than attentional filtering.

**Impact Statement:** We found that when visual stimuli are paired with sounds, they can induce fluctuating activity in neurons that are primarily sensitive to sound alone. Such activity fluctuations may permit the representation of both the visual and auditory stimuli in an overlapping population of neurons.

## Introduction

How the brain encodes multiple events in the environment is not well understood. A recent theory suggests that encoding multiple stimuli may at least partially involve neurons switching back and forth between responding to different items across time, or multiplexing (Caruso et al., 2018; Groh et al., 2024). To date, empirical work concerning multiplexing has focused on stimuli within a single sensory modality – two visual stimuli (Caruso et al., 2018; N. Y. Jun et al., 2022; Schmehl et al., 2024; Chen et al., 2024) or two sounds (Caruso et al., 2018; Chen et al., 2024). The rationale for this has been to study cases in which stimuli can be thought of as competing for representation within the same general population of neurons. For example, two drifting gratings, positioned adjacent to one another and simply drifting in opposite directions, are similar in kind and equally capable of evoking responses in areas such as V1, MT, and V4. Similarly, two bandpass sounds, differing only in frequency and location, are equally capable of evoking responses in the inferior colliculus (IC). Arguably, such stimuli have equal “standing” with regard to the neural population under study, and these cases have been shown to evoke multiplexing responses in some neurons in these four brain regions (Caruso et al., 2018; N. Y. Jun et al., 2022; Schmehl et al., 2024; Chen et al., 2024).

In contrast to these situations involving stimuli that are both similarly “apt” for a given brain region, multiplexing has also been identified when only one of two stimuli is well-suited to the brain area under study. For example, the face patches of inferotemporal (IT) cortex are so named because they consist of neurons that are selective for face stimuli. These neurons are more sensitive to faces compared to non-face objects and even show selectivity to specific faces. Intriguingly, when we tested responses in IT cortex to combinations of two faces as well as combinations of a face and a non-face object, we found multiplexing responses for both face-face and face-object combinations, albeit to different degrees in different face patches (MF vs. AL) (Schmehl et al., 2024). These findings raise the question of how selectivity for a particular class of stimuli impacts the presence and prevalence of switching patterns.

An ideal case for exploring this question involves combinations of stimuli of different sensory modalities, such as visual and auditory stimuli – which are initially detected by completely distinct sense organs (eyes vs. ears). While emerging evidence highlights that there is crosstalk between sensory systems at a variety of levels of the pathway (Bulkin & Groh, 2006; Ghazanfar & Schroeder, 2006; Kayser & Logothetis, 2007; Gruters & Groh, 2012; Teichert & Bolz, 2018; Schmehl & Groh, 2021), and descending projections can carry cross-sensory signals to very early points including the sensory periphery (Gruters et al., 2018; Lovich et al., 2023; Bröhl & Kayser, 2023; Sotero Silva et al., 2025; Abbasi et al., 2025), it remains the case that some areas of the brain are far more sensitive to one sensory modality than another. Does multiplexing occur in such predominantly unimodal areas even when one of the two stimuli being presented is in the non-dominant modality?

Here, we investigate that question by assessing the responses to combinations of visual and auditory stimuli in the primate inferior colliculus, a predominantly unimodal auditory structure located only a few synapses in from the ear itself (for review, see (Winer & Schreiner, 2005)). The IC is known to be responsive to visual as well as auditory stimuli, although visual stimuli evoke much weaker changes in firing than auditory stimuli do (Porter et al., 2007; Bulkin & Groh, 2012; Schmehl et al., 2025). As noted above, some IC neurons exhibit fluctuating activity in response to two sounds (Caruso et al., 2018). Here, we ask several questions about the effect of visual stimuli on the IC’s fluctuating patterns. First, do such fluctuations also occur when the stimuli consist of a sound and a visual stimulus? If so, this would suggest that the IC keeps visual and auditory signals distinct, potentially preserving information about both. Second, we turn to what happens when a visual stimulus accompanies one of two sounds – does the visual stimulus cause IC neurons to preferentially encode the sound it is paired with? Adding a salient visual cue to one of the sounds might bias the representation to favor one sound over the other.

To investigate these questions, we recorded IC single-unit activity while monkeys performed a saccade task involving visual, auditory, and/or combined audiovisual stimuli presented at one or two locations. We found fluctuating activity evoked by audiovisual stimuli in some IC neurons, suggesting that even when a sound and visual stimulus co-occur at the same location, IC neurons maintain distinct signals about both. When two sounds were presented at different locations, fluctuating activity occurred regardless of whether any of the sounds were paired with a visual stimulus. In contrast with related results in auditory cortex (Atilgan et al., 2018), the encoding of sound was not generally biased toward the visually-paired sound. Both sounds were represented even though the task required the monkey to report them sequentially. Indeed, in one monkey, the IC showed a bias favoring representation of the *second* location that they made a saccade to. Together, these experiments suggest that the IC retains information about both auditory and visual stimuli, supporting flexible behavioral performance.

## Materials and Methods

### Subjects and Surgical Preparation

Here we report results from the same experiments reported in (Schmehl et al., 2025), now also including additional conditions not reported previously. Two rhesus macaques (*Macaca mulatta*, one female age 13 years and one male age 4 years) participated in the study. All procedures were approved by the Duke University Institutional Animal Care and Use Committee and were conducted at Duke University. Monkeys were prepared for experiments by implanting a head restraint device and a recording cylinder (Crist Instruments) via two stereotactic surgeries under isoflurane anesthesia. Surgeries were conducted using aseptic techniques and appropriate post-surgical analgesics. The placement of the recording cylinder allowed the inferior colliculi in both hemispheres to be accessible at an angle of 26 degrees from vertical in the coronal plane. A coronal magnetic resonance imaging scan confirmed cylinder placement. Electrodes were held in placed by a plastic grid in the recording cylinder (Crist Instruments).

### Equipment, Stimuli, and Task

Monkeys were trained to perform a localization task involving combinations of sounds and visual stimuli (Figure 2a). Stimulus presentation and eye position monitoring (video eye tracking, 500 Hz sampling) were controlled using custom software (“Beethoven” by Ryklin Software, Inc.). Sounds were presented from speakers (Cambridge SoundWorks) and consisted of bandpass filtered noise at center frequencies of 420 or 2000 Hz (11 kHz sampling rate, 55 dB SPL, ±100 Hz bandwidth, 10 ms on ramp). Waveforms were computed once for each recording session and were thus the same across trials for a given sound frequency for a given neuron. They were calibrated to have the same sound level across speakers. Visual stimuli consisted of light-emitting diodes (LEDs) attached to the fronts of the speakers. All experiments were conducted in a dark single-walled sound attenuation chamber (Industrial Acoustics Company, lined with Sonex sound attenuating foam).

Each trial began with the onset of a central fixation light located straight ahead (i.e., on the midline and at eye level). After the monkey fixated for 600-700 ms, a target condition was presented involving stimuli at either one or two locations, 12 degrees left or right on the horizontal meridian. Stimuli could involve sounds alone, visual stimuli alone, or combinations of visual and auditory stimuli. On single-location trials, only one location was used; the stimulus could be auditory, visual, or combined audiovisual at the same location. Detailed results from these single-location trials are reported in (Schmehl et al., 2025). On two-location trials, both locations were used, and these could involve either two sounds, a sound and a sound-light combination, or two sound-light combinations. Regardless, the monkey maintained fixation on the initial fixation light until the fixation light was extinguished (600-800 ms after target onset), and subsequently made saccades to all presented target locations, with either order accepted as correct on the two-location trials. Trials of all types were judged correct if the monkey made saccades to each presented target location within a window of 16 degrees horizontally by 60 degrees vertically and fixated there for 200-250 ms, after which a juice reward was delivered. These reinforcement windows were larger than needed for visual-only trials, but allowed for less precision on auditory trials, particularly in the vertical dimension (Jay & Sparks, 1990; see also: Gnadt et al., 1991; Barton & Sparks, 2001 for consideration of saccade accuracy in non-visual tasks). Conditions were randomly interleaved and were weighted evenly for one-location vs. two-location conditions. Conditions of each modality were weighted 30% visual, 30% auditory, and 40% audiovisual.

### Neural Recordings

Single-unit extracellular recordings were conducted in the inferior colliculus using a 125 mm tungsten microelectrode (FHC) held in place via a plastic grid inside a recording chamber (Crist Instruments, grid hole spacing 1 mm). Spiking activity was collected with a Multichannel Acquisition Processor (MAP system, Plexon Inc.), passed through appropriate amplifiers, and sent to SortClient software for online spike sorting. Single unit spiking activity was identified from signals that were bandpass filtered between 150 Hz and 8 kHz and sampled at 20 kHz. Spike times, eye positions, and task parameters were also stored for offline analysis.

As described previously (Schmehl et al., 2025), we recorded from 106 sound-responsive neurons, as assessed via visual inspection of the peristimulus time histograms and/or statistically during a transient (0-100 ms) or sustained (0-500 ms) response window in comparison to a corresponding pre-stimulus baseline period (paired t-test, p<0.025 for each tail). At least 250 correct behavioral trials were collected for each of these neurons. 18 were also responsive to visual stimuli (paired two-tailed t-test comparing activity 50-250 ms after stimulus onset to an equivalent baseline period, p<0.025 for each tail). 82 were from the right IC (Monkey J), and 24 were from the left IC (Monkey D).

### Analysis of fluctuating activity

#### Model 1: Involving Benchmarked Poissons and Mixtures of Poissons

To determine how neurons respond to two simultaneous targets, we used statistical techniques previously developed by the Groh and Tokdar labs (Caruso et al., 2018; Mohl et al., 2020; Glynn et al., 2021). The first stage of analysis compares neural responses from trial to trial, describing a neuron’s response on dual-stimulus trials (AB) in relation to the response distributions on trials when each stimulus was presented individually (A or B). We modeled the trial-wise spike count distributions in response to targets A and B as Poisson distributions and asked how the AB spike count distributions compared to the A and B distributions. We analyzed “triplets” of one A condition, one B condition, and one AB condition, and categorized each triplet into one of four labels based on the label with the highest posterior probability after a Bayesian model comparison beginning with equal priors (Figure 1a):

1. *Outside*: The AB response distribution is Poisson with a mean that is either greater than the higher of the means of the A and B responses, indicating a facilitative combination of the A and B signals, or lower than the lesser of the means of the A and B responses, indicating a suppressive combination of those signals.
2. *Single*: The AB response distribution is Poisson with a mean matching that of either A or B, indicating a winner-take-all (or loser-take-all) response.
3. *Intermediate*: The AB response distribution is Poisson with a mean between the A and B means, indicating either averaging of the A and B responses or signals fluctuating between A and B at a rapid time scale.
4. *Mixture*: The AB response distribution is drawn from both the A and B distributions, creating a bimodal AB response distribution with peaks centered at the A and B means. This response pattern indicates alternation between the A and B responses from trial to trial.

**Figure 1:**
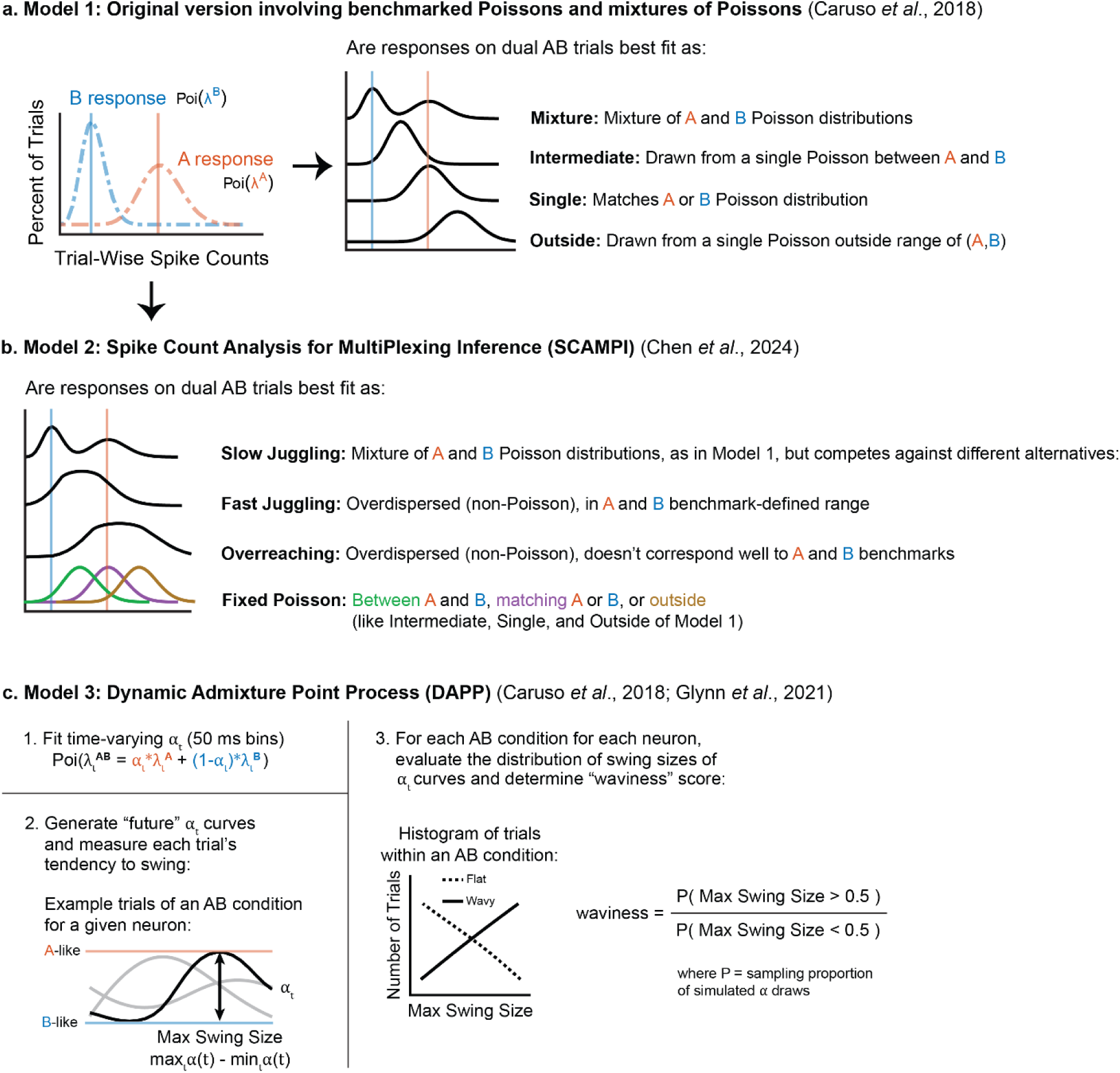
Modeling approaches for evaluating responses to combinations of stimuli. Conditions are assigned into “triplets” consisting of one condition where stimulus A was presented (red), one condition where stimulus B was presented (blue), and one condition where both were presented (AB, black curves). We assume trial-wise (Models 1 and 2) or bin-wise (Model 3) spike counts for single stimuli follow a Poisson distribution. a) In the first model (Caruso et al., 2018; Mohl et al., 2020; N. Y. Jun et al., 2022; Schmehl et al., 2024; Groh et al., 2024), the AB spike count distribution may be: a *Mixture* of the A and B distributions, suggesting trial-to-trial fluctuations between the responses to the two single stimuli; a Poisson distribution with an *Intermediate* mean between the A and B means, suggesting averaging (or very even amounts of fluctuation within trials – see Model 2); a Poisson distribution with a mean rate that matches either the *Single* A or B mean, suggesting a winner-take-all strategy; or a Poisson distribution with mean rate *Outside* the range bounded by the A and B means, suggesting addition or subtraction of the responses. b) In the second model (Chen et al., 2024), new categories are added to allow for additional overdispersed-Poisson alternatives, and the three Poisson categories from Model 1 (Intermediate, Single, and Outside) are here grouped into a single option (Fixed). The overdispersed-Poisson options consist of *Slow Juggling*, corresponding to across-trial fluctuations, *Fast Juggling*, corresponding to fluctuations that may occur at least partially within trials, but too unevenly to produce a single Poisson distribution, and Overreaching, which indicates that activity is fluctuating but that it is not well captured by the A- or B-distribution benchmarks. c) Finally, we deploy the Dynamic Admixture Point Process model (Caruso et al., 2018; Glynn et al., 2021) to quantify how much fluctuation can be observed within individual trials. This model operates at the subtrial scale, here spike counts in 50 ms bins across time in each trial, and fits the AB sequence of bin-wise spike counts as a weighted admixture of those typically observed on the A-alone and B-alone trials. The observed weight curves are used to generate predicted weight curves for hypothetical future trials. A “waviness” metric is computed on these future weight curves.

Unless otherwise noted, in this analysis, data for each trial was analyzed during the period between 0-500 ms after stimulus onset, before the eyes have moved.

#### Model 2: SCAMPI – a Model With Expanded Non-Poisson Alternatives

To evaluate response patterns against a greater range of non-Poisson options, we used a revised analysis model, known as the Spike Count Analysis for MultiPlexing Inference (SCAMPI) model (Chen et al., 2024). This model permits the detection of fluctuation with an altered set of classification categories, setting up different forms of competition and providing more flexibility with regard to the role of the A and B response benchmarks (Figure 1b):

1. *Slow Juggling*: The AB response distribution is drawn from both the A and B distributions, creating a bimodal AB response distribution with peaks centered at the A and B means. This response pattern is logically analogous to the first model’s *Mixture* category, indicating alternation between the A and B responses from trial to trial, but differs from the first model in now having to compete against several alternatives that are also not based on a single Poisson distribution.
2. *Fast Juggling*: The AB response distribution is overdispersed relative to a single Poisson, but still bounded by the A and B distributions. This response pattern indicates more rapid fluctuation between the A and B responses, with one or more switches happening within the course of individual trials. The within-trial switches cause additional trial-to-trial spike count variability, which should manifest as overdispersion but may not lead to bimodality as in the earlier case of Slow Juggling.
3. *Overreaching*: The AB response distribution is overdispersed relative to a single Poisson but not well bounded by the A and B benchmarks. This response pattern indicates that fluctuations may be occurring, but that they are not necessarily fully explained as switching between the benchmark A and B response means.
4. *Fixed*: The AB response distribution is a single Poisson distribution, indicating no fluctuating activity above and beyond that intrinsically captured under a Poisson process. The mean firing rate could match one of the benchmarks (like Single), be between them (like Intermediate), or be outside the range (like Outside) – all such cases are considered “Fixed” so long as the distribution of spike counts is well described as a single Poisson.

#### Model 3: Dynamic Admixture Point Process (DAPP) Model

The SCAMPI model (Model 2) permits detecting fluctuations at fast time scales, i.e., when switching intervals are shorter than the length of a single trial. A detailed picture of such rapid fluctuations can be constructed by analyzing the underlying spike trains with the Dynamic Admixture Point Process (DAPP) Model (Caruso et al., 2018; Glynn et al., 2021) (Model 3, Figure 1c). The DAPP Model is a within-trial analysis method that treats spiking activity on AB trials as an admixture of the A and B spiking activity. The model operates on spike counts calculated in a series of bins (here, 50 ms bins, in a sequence of 10 for any given trial), and evaluates the tendency of the activity to “swing” back and forth between the A and B rates for each bin over the course of the trial. The model can predict hypothetical future AB trials and quantify the typical “swing size” of any fluctuations as the response “swings” between A and B. A score for “waviness” is calculated as the odds of the response swinging at least 50% between its peak and trough, where a higher waviness value indicates a greater degree of fluctuation between the A and B responses during individual trials.

## Results

### Behavior

We trained two monkeys to localize sounds, LED lights, and sound-light pairs arising from one or two simultaneous locations (Figure 2a). Sounds were bandpass noise at center frequencies of 420 or 2000 Hz, counterbalanced for the two locations (−12 or +12 horizontal degrees) and for the presence or absence of a paired light. Monkeys made eye movements to the stimulus locations on interleaved one-location and two-location trials. Both monkeys performed both types of trials correctly. Monkey D performed equally well on both the one- and the two-location trials, (82.3 and 80.7% correct) whereas Monkey J was slightly better at the one-location trials (84.8% correct for one-location, 77.8% correct for two-location) (Figure 2b,c).

**Figure 2:**
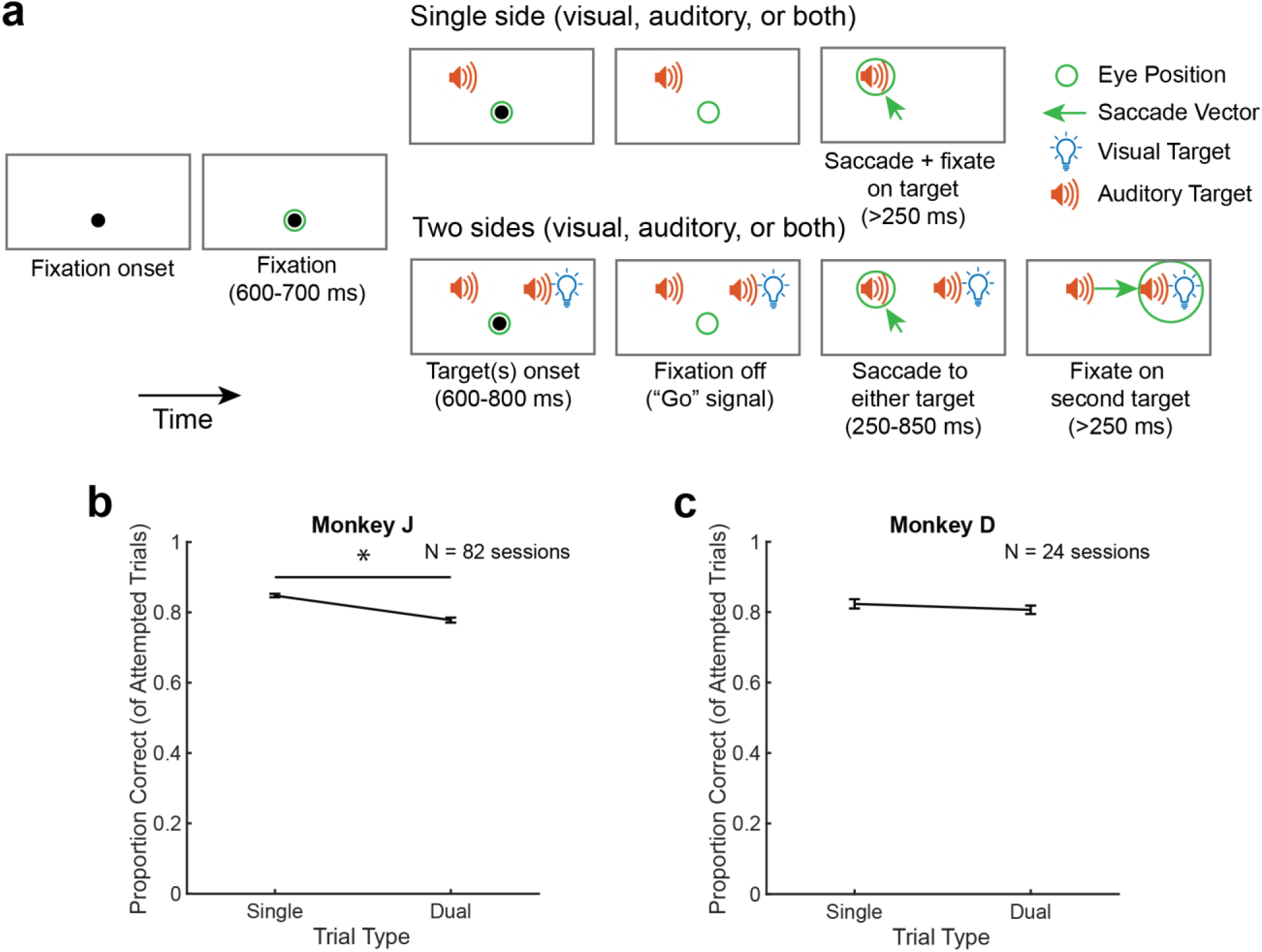
Monkeys accurately perform a two-location audiovisual localization task. a) Task structure. Each trial began with the onset of a central light, and the monkey initiated the trial by fixating on the light for a variable period between 600 and 700 ms. The target(s) then turned on, and the monkey waited 600-800 ms for the central fixation light to disappear, cuing it to make a saccade to each target (green arrows). If two targets were present, the monkey could localize the targets in either order. After localizing all targets in the trial, the monkey received a juice reward. b-c) Behavioral performance for Monkey J (b) and Monkey D (c). Performance grouped by number of targets (single, one target; dual, two targets). Proportion correct is calculated based on attempted trials, defined as those in which the monkey initiated the trial by fixating for the appropriate time period. Percent correct was calculated within sessions and results are shown as the average percent correct across sessions, with error bars indicating standard error across sessions. Chance performance is not straightforward to compute for spatially and temporally continuous behavior, but could be roughly estimated as 20% for the single location trials (16 degree reinforcement window out of an 80 degree oculomotor range in the horizontal dimension). Chance for dual location trials would be less than this because the eyes had to enter and pause in both reinforcement windows.

### Neural responses to combined visual and auditory stimuli at the same location

We previously showed that the *average* responses to audiovisual stimuli differed statistically from the auditory responses in only about 23.5% of the conditions tested in this dataset of IC neurons (Schmehl et al., 2025). Adding a visual cue to a sound at the same location could either slightly increase (6.8%) or slightly decrease (16.7%) the activity compared to when the sound was presented alone (Schmehl et al., 2025). However, since these observations were based on the average activity across trials, it is unknown whether *neurons might exhibit fluctuating activity, potentially encoding the visual stimulus some of the time and the sound the rest of the time*. If so, this would indicate that the neurons in this brain region are sensitive to the presence of the visual stimuli, even though their average firing rate may not be statistically different between sound-only and audiovisual stimulus presentation as ascertained using conventional methods such as t-tests as we did in our previous paper (Schmehl et al., 2025). Accordingly, we investigated activity fluctuations in the entire dataset, not limiting ourselves to cases in which the average firing rate on audiovisual trials was statistically distinguishable from the corresponding sound-alone trials.

We first used our previously developed statistical method for assessing fluctuating activity (Caruso et al., 2018; Mohl et al., 2020). In this analysis method, a neuron’s response to two simultaneous stimuli (AB, audiovisual) is evaluated in relation to the response distributions when each target was presented individually (A or B, auditory or visual alone). “Triplets” of one A condition, one B condition, and one AB condition were classified into one of four categories (Model 1, Figure 1a). The main category we are interested in is the “Mixture” category, in which the spike counts on AB trials appear to be drawn from a mixture of the A-only and B-only spike count distributions. “Mixtures” compete against three other categories: “Single,” in which the spike counts on AB trials appear to be consistently drawn from either the A or the B distributions, but not both; “Outside,” in which the spike counts on AB trials appear to be outside the range of spike counts observed on the A and B trials, as would occur if the neuron responded more strongly on the AB trials than on either the A or B-alone trials; or “Intermediate,” in which the spike counts on AB trials appeared to be drawn from a consistent distribution with a mean between that of the A-alone and B-alone spike count distributions. Note that “Intermediate” is an ambiguous category that could arise from activity fluctuations that were too fast to be observed in the comparatively long spike counting windows (500 ms) analyzed here.

While this analysis method is not limited to cases involving a statistically significant difference between the audiovisual and auditory response conditions, it nevertheless cannot be run on every triplet – two additional conditions must be met. First, the responses to A and B alone must differ from one another enough that it is possible to tell the difference between the potential outcomes (Mixture, Single, and Intermediate would not meaningfully differ if the responses to A and B were highly similar). We assessed the likelihood that the A and B responses were drawn from the same vs. a different distribution, and included only cases in which A and B trials being drawn from different distributions was substantially more likely than being drawn from the same distribution (Bayes factor > 20). A total of 229 of 413 potential triplets survived this screening. The overwhelming majority of such triplets exhibited excitation to the auditory stimulus and only a very minimal change in activity evoked by the visual stimulus, consistent with our prior report concerning this dataset (Schmehl et al., 2025). Figure 3a shows the distribution of responses, expressed as Z-scores in relation to a pre-stimulus baseline. Sounds evoked a wide range of response patterns ranging from mild suppression (Z=-2.4 standard deviation units relative to baseline) to strong excitation (Z=60.8), with a median value of Z=2.23, whereas the direct impact of visual stimuli was much more modest (Z=-1.7 to 1.80, median −0.03). Figure 3b shows the differences between the auditory and visual response Z-scores within triplets; the vast majority involve stronger auditory than visual responses.

**Figure 3:**
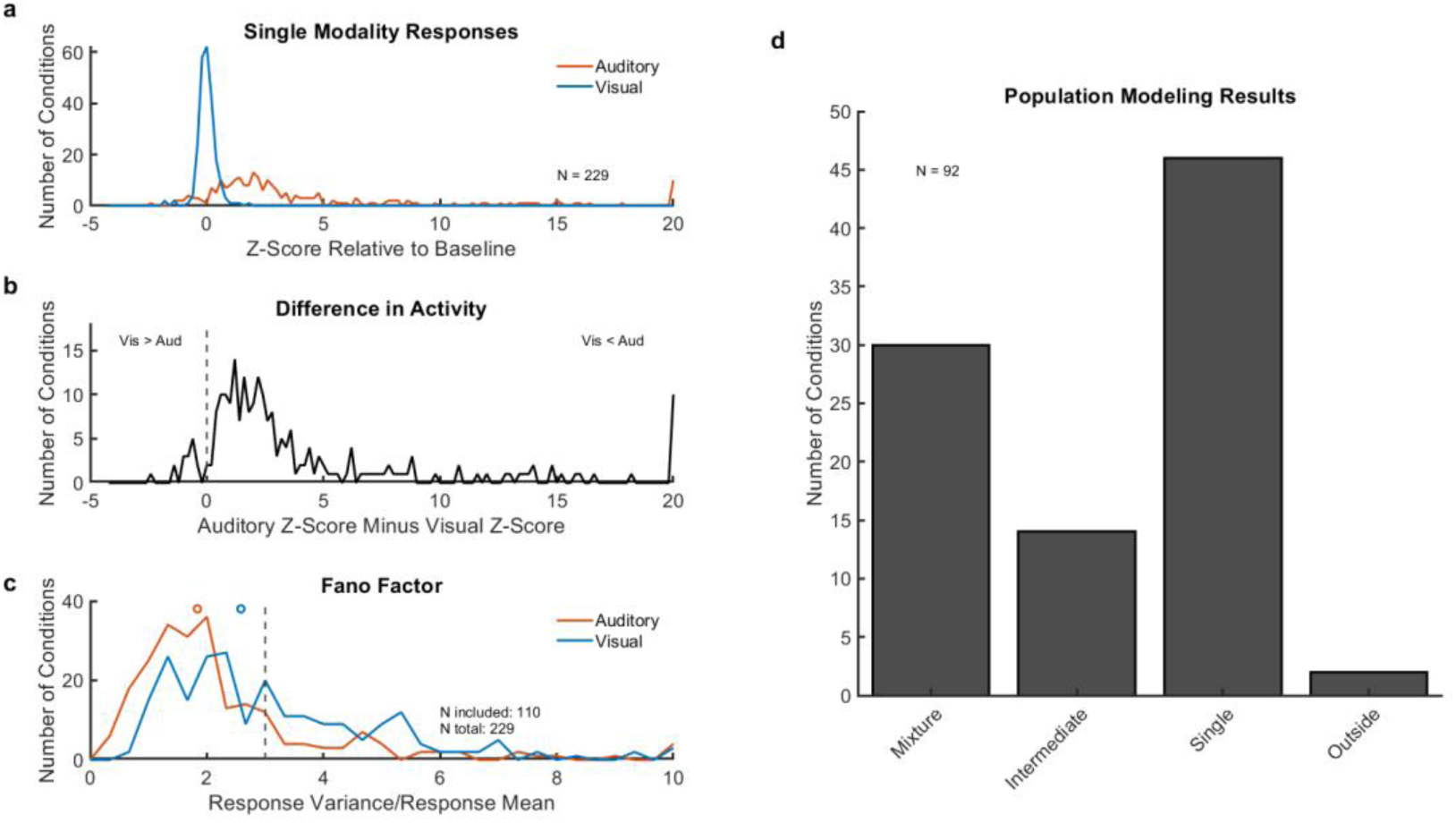
Screening and modeling results for a sound-light pair. (a-b). Triplets could only be included if there was a sufficient difference in the mean response on auditory-alone and visual-alone trials. Panel (a) shows the distribution of mean responses on the auditory (red) and visual (blue) conditions, expressed as a Z-score relative to a matching baseline window of activity (baseline: −500 to 0 ms relative to stimulus onset vs. sensory: 0-500 ms after stimulus onset). Panel (b) shows the difference between these two mean Z-scores for each triplet. These panels show the N=229 triplets that showed sufficiently different levels of activity for potential inclusion in the modeling analysis. (c) In addition to exhibiting a difference in the auditory-alone and visual-alone responses, triplets were only included if the variance-to-mean ratio (Fano factor) of the spike count responses on auditory-only (red) and visual-only (blue) trials were reasonably well characterized as following a Poisson distribution (Fano ∼ 1). Open circles indicate the median values; dashed line illustrates the inclusion criteria (Fano < 3). After application of both response-difference and Fano factor screening, 110 triplets were included for further analysis. d) Results of the trial-to-trial analysis for triplets classified with a probability greater than 50% (N=92). Half of these triplets are classified as Single, all of which responded only to the sound. About a third are classified as Mixtures, indicating a high degree of trial-to-trial fluctuation between the single auditory and single visual responses. See also Supplementary Figure 1 for analysis of a 100-500 ms spike counting time window.

The second inclusion criterion is that the A and B response distributions each had to be reasonably well described by a Poisson distribution; if this is not true, then the assumptions of the statistical analysis would be violated and the results could be inaccurate. To exclude cases violating the Poisson assumption in this dataset, we screened out triplets with high variance-to-mean ratio of the A or the B responses. For a Poisson distribution, the true variance-to-mean ratio – known as the Fano factor – should equal 1. Figure 3c shows the distribution of Fano factors for the auditory-only (red) and visual-only (blue) conditions in the dataset (after passing the screen for differing auditory vs. visual responses, as described above). Generally, the Fano factors for the auditory-only trials were roughly consistent with a Poisson distribution, although a bit more variable than expected, with a median Fano=1.83 (red circle). The Fano factors for the visual-only trials were higher than for the auditory trials, with a median of 2.58 (blue circle). The reasons for this difference are uncertain. Regardless, following our previous studies (Schmehl et al., 2024; Chen et al., 2024), we selected triplets for which both A and B Fano factors were less than 3. All analyses were also conducted with more strict Fano-screening criteria, i.e., both A and B response patterns having a Fano factor less than 2, and results were similar but involved a smaller number of included conditions.

Figure 3d shows the results of the trial-to-trial analysis comparing responses to a sound, a light, and a sound-light pair occurring at the same location in space. Not every triplet could be well characterized with high confidence, but among those for which the winning model was more likely than all the other models combined (winning probability > 0.5), half were classified as Single (46 of 92, or 50.0%) – all of which responded chiefly to the auditory stimulus and appeared to be unaffected by the visual stimulus. Note that this is smaller than the roughly 75% that failed to show a statistically significant difference in the average response between auditory and audiovisual trials on the basis of t-tests (Schmehl et al., 2025). Indeed, a sizeable fraction of the dataset (30 of 92, or 32.6%) were classified as Mixtures, indicating fluctuation between the response to the sound and the response to the visual cue on individual trials (Figure 3d). Such fluctuating activity is likely to be overlooked in conventional statistical methods, which examine average activity across many trials.

A possible confound in this analysis is that the latency of auditory responses is shorter than for visual responses. Thus, responses early in this 0-500 ms spike counting window might be dominated by the sound alone for trivial reasons. To rule out this possibility, we repeated the analysis on a time window of 100-500 ms (beginning after the latest visual latency in this dataset), in comparison to a baseline period of equivalent duration (Supplementary Figure 1). The proportion of Mixtures was similar: 31 of 81 or 38.3% of conditions were classified as Mixtures, i.e., slightly higher than for the 0-500 ms spike counting window. This shows that fluctuation between the auditory and visual responses is robust to the time window of analysis and doesn’t appear to be a consequence of the visual response arising later in the trial than the auditory response.

Examples of Mixture responses from individual neurons responding to individual conditions are shown in Figures 4 and 5. In each of these examples, the distribution of responses on audiovisual trials (gray bars: histogram of individual trials; black lines: spline fit) showed evidence of being drawn from a mixture of the distributions observed on corresponding auditory-alone and visual-alone trials. In some of these examples (Figure 4), the fluctuating activity is well matched to the corresponding single-modality trials, whereas in others (Figure 5), the distribution of trials is less clearly related to the underlying A and B benchmarks, but they were nevertheless classified as Mixtures.

**Figure 4:**
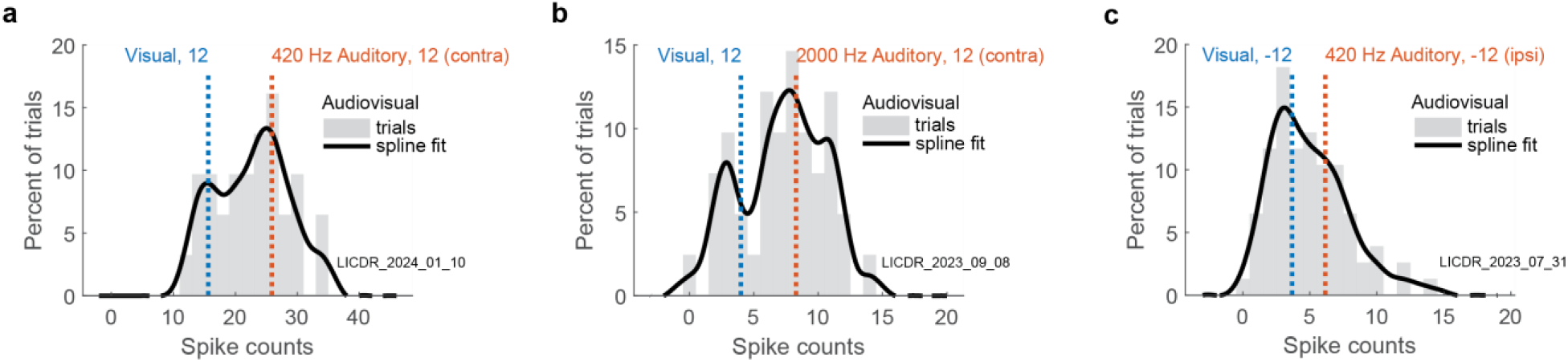
Fluctuating activity preserves information about an individual sound and light in a sound-light pair. Example neurons’ Mixture responses to individual conditions consisting of one sound, one light, and one sound-light pair, all at the same location in space. Black solid lines represent sound-light pair responses, and red and blue dotted lines represent mean auditory and visual responses, respectively. Sound-light pair responses were smoothed via convolution with a 3-point sliding window, with the middle point doubly weighted compared to the first and last points, with cubic spline interpolation. Raw distributions are depicted in gray bars. Spikes were counted in a window of 500 ms following stimulus onset. In all three examples, sound-light pair responses exhibit modes that roughly correspond to the single auditory and visual means.

**Figure 5:**
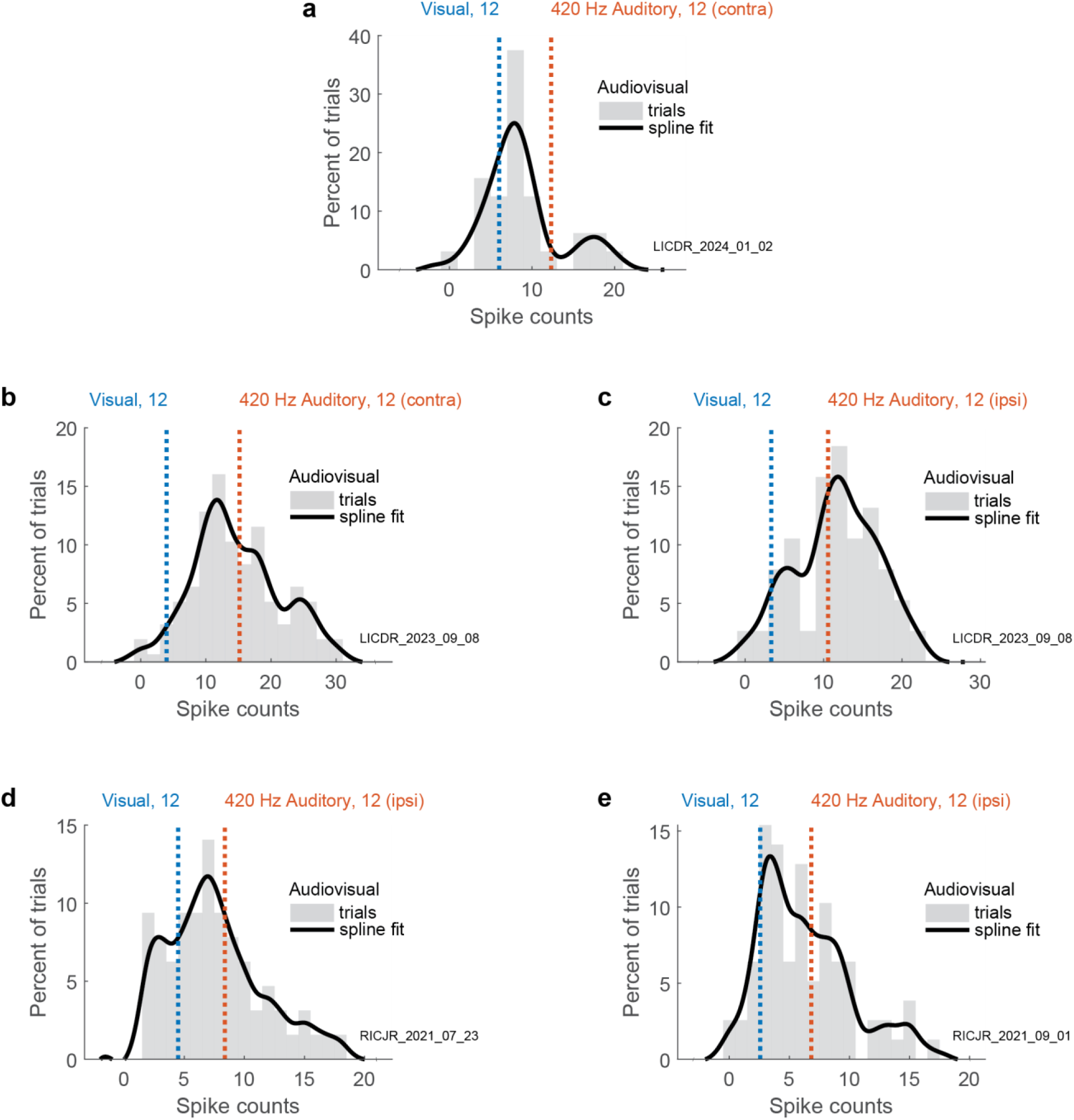
Fluctuating activity may not always be precisely related to benchmarks defined by an individual sound and light. Same as Figure 4, but for Mixture triplets whose sound-light pair responses have modes that appear not to be well-benchmarked by the single auditory and visual means.

To further evaluate this pattern in relation to the benchmarking, we conducted an additional round of modeling on the Mixture category, using a second modeling approach involving an expanded range of non-Poisson alternatives (Figure 1b). Like the first model, this model (called “SCAMPI,” or Spike Count Analysis for MultiPlexing Inference, (Chen et al., 2024)) evaluates the spike count distributions on dual-stimulus trials in relation to four competing possibilities, but it differs in two key respects. First, it includes multiple over-dispersed (non-Poisson) alternatives. Second, it allows for the possibility that the single-stimulus benchmarks do not provide a good account for the dual-stimulus response patterns (Model 2, Figure 1). Specifically, this modeling approach offers three outcomes that are overdispersed compared to a true Poisson, differing from each other in the presumed underlying rate of switching and whether they are accurately captured by the single stimulus benchmarks. The “Slow Juggling” category is a strict category in which it is presumed that a given neuron responds to one and only one stimulus throughout the spike counting window, and that which stimulus will be favored for a particular trial is determined by some process that takes place between trials. The “Fast Juggling” category is more forgiving. It assumes that while at any given moment a neuron is responding to only one of the two stimuli, the neuron may switch from one to the other during the course of a stimulus presentation, leading to a distribution of spike counts that is overdispersed relative to a Poisson distribution, but still bounded by the single-stimulus benchmarks. This category may still show clear modes near these benchmarks, but they may be less distinct than is expected of Slow Juggling. The third non-Poisson outcome is one in which the distribution of spike counts is overdispersed, but does not seem to be clearly constrained by the single-stimulus benchmarks (“Overreaching”). These three outcome categories must each compete against a comprehensive Fixed Poisson distribution category, which can have any relationship to the single stimulus benchmarks, thus encompassing the Intermediate, Single, and Outside categories from Model 1.

We ran this second model on the conditions categorized as Mixtures in the first model (detailed categorization shown in Supplementary Figure 2). 28 of 30 triplets were confirmed as having non-Poisson distributions, of which 9 were well characterized as constrained by the benchmarks (1 Slow, 8 Fast) and 19 were classified as overdispersed but not fully constrained by the benchmarks (Overreaching). The examples shown in Figure 4 were classified as Fast, and the examples shown in Figure 5 were classified as Overreaching. This analysis supports the interpretation that the addition of a visual stimulus to an auditory stimulus can induce patterns of fluctuating activity, but that the fluctuating activity is not always exactly captured by the single-stimulus benchmarks. We will return to the interpretation of the potentially shifting single-stimulus benchmarks in the Discussion.

Overall, these results indicate that fluctuating activity is evident in the IC in this multisensory context. Specifically, they suggest that information about the sound alone is still present in the neural firing patterns, even when accompanied by a visual stimulus. Furthermore, this finding extends prior reports of fluctuating activity elicited by stimuli at two different locations (Caruso et al., 2018; N. Y. Jun et al., 2022; Schmehl et al., 2024; Chen et al., 2024) to a case where both stimuli arise at the same location – thus suggesting that shifting spatial attention cannot fully account for the presence of such activity patterns. We will return to this topic in the Discussion as well.

### Influence of visual stimuli on encoding of two sounds at different locations

We next extended this approach to explore how the IC responds when two sounds are presented at different locations, and a visual stimulus is paired with one of them. Perceptually, a paired visual stimulus can boost the salience of a sound, or even capture its perceived location (Thomas, 1941; Jackson, 1953; Thurlow & Jack, 1973; Bertelson & Radeau, 1981; Alais & Burr, 2004; Atilgan et al., 2018), but it is unclear what the underlying neural basis of this phenomenon might be (Callan et al., 2015; Bruns, 2019). We have previously shown that some neurons in the IC fluctuate when two sounds are present (Caruso et al., 2018). Here, we explored whether the presence of a visual stimulus paired with one of these two sounds would reduce the presence of such fluctuating signals, potentially in favor of the visually-paired sound being over-represented in comparison to the sound-only stimulus.

We first verified that two sounds presented without visual stimuli yielded fluctuating activity patterns in this dataset. Using the same modeling analysis and inclusion criteria described above, we tested the response patterns to triplets consisting of a 420 Hz bandpass sound at one location, a 2000 Hz bandpass sound at the other location, and both sounds simultaneously. Figure 6 shows the results. Generally, the lower frequency sound (420 Hz) elicited stronger responses than the higher frequency sound, consistent with the over-representation of low frequency sounds in the primate IC (Figure 6a,b); only about 17-18% of the IC is thought to be tonotopic and thus to include selectivity for high frequency sounds – the remainder is either selective for low frequencies or insensitive to frequency (Bulkin & Groh, 2011; Ress & Chandrasekaran, 2013). However, there was little difference in the Fano factors (low frequency median Fano: 1.7, high frequency median Fano: 2.0) (Figure 6c). A total of 41 triplets survived both response-difference and Fano screening, of which 34 produced a model winner with a winning probability greater than 0.5 (Figure 6d). Of these, Single was the most common winner (14 of 34, or 41.1%), and of these, 12 involved the low frequency sound “winning” (85.7%), and 2 involved the high frequency sound “winning” (14.3%). Mixtures occurred in 8 of 34, or 23.5% of conditions, which was somewhat less common than on the audiovisual conditions described in Figure 3. As with the audiovisual conditions, auditory-auditory Mixtures contained both Fast Juggling (2 of 8, or 25%) and Overreaching cases (6 of 8, or 75%), suggesting that the benchmarks do not always precisely demarcate the fluctuating activity patterns. However, in net, between the steady responses of the Single triplets and the fluctuating patterns observed in the Mixture triplets, the inference is that both sounds were represented in the IC, even if not necessarily weighted equally in this sample.

**Figure 6:**
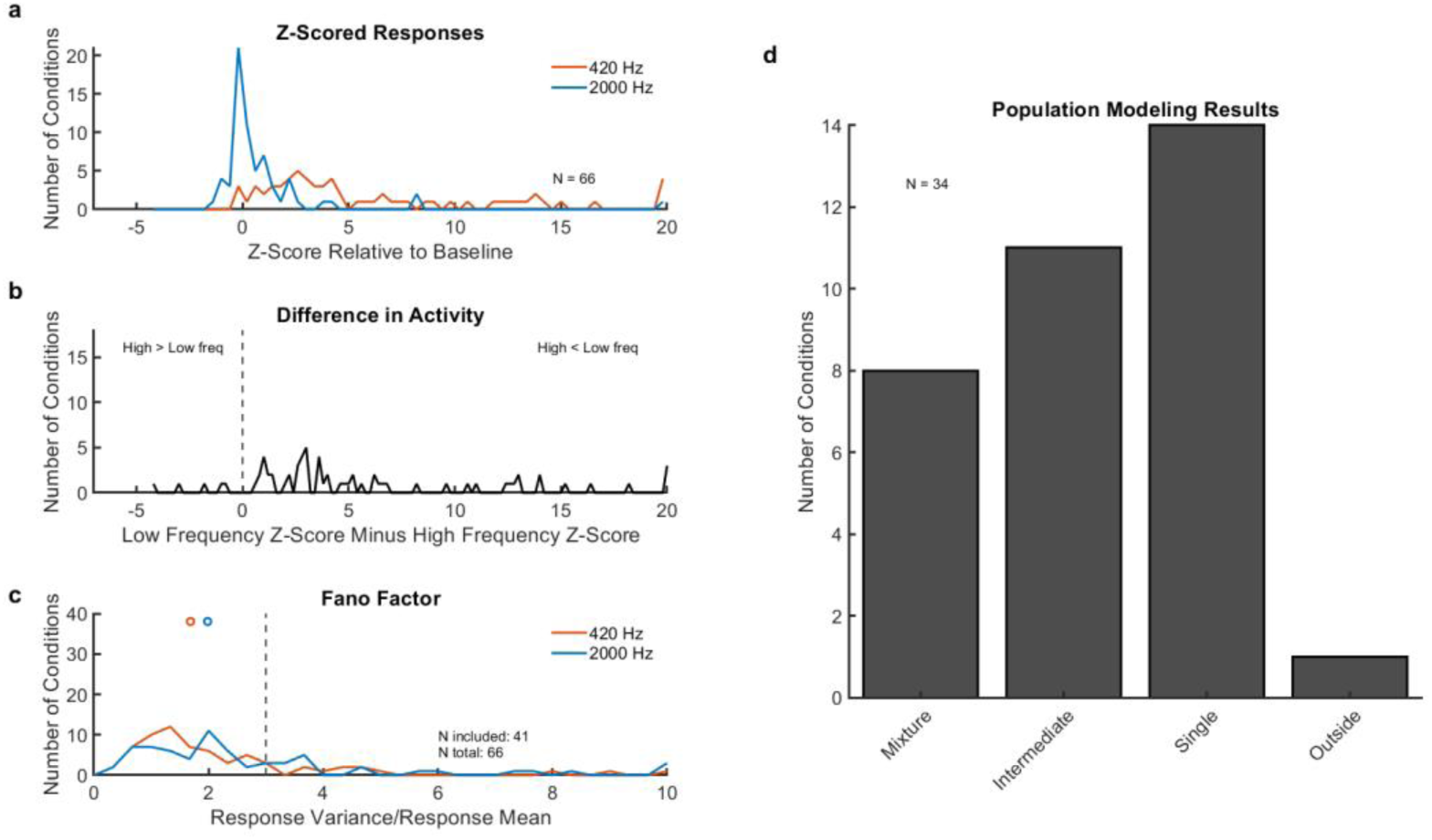
Screening and modeling results for two simultaneous sounds. In all panels, analysis and plotting conventions are the same as for Figure 3. a) Distribution of mean responses on the 420 Hz (red) and 2000 Hz (blue) conditions. b) Difference between these two mean Z-scores for each triplet. N=66 showed sufficiently different levels of activity for inclusion. c) Variance-to-mean ratio (Fano factor) of the spike count responses on 420 Hz (red) and 2000 Hz (blue) conditions. 41 triplets were included for further analysis. d) Results of the trial-to-trial analysis for triplets classified with a probability greater than 50%. Singles and Mixtures collectively make up 64.6% of conditions, indicating that information is preserved about both sounds across neurons (Singles) and from trial to trial within neurons (Mixtures).

Having verified the presence of fluctuations for two sounds, we next turned to the trials involving two sounds and a visual stimulus paired with one of them. Triplets were constructed from trials involving a single sound at one location (A), an audiovisual combination at the other location (dubbed B, although note that they consist of two stimuli together), and all three together (AB). We continue to use the “triplet” terminology because there are still three groups of trials to be analyzed, of which two serve as benchmarks for analyzing the third.

Figure 7a-c shows the response difference and Fano factor screening results. The mean audiovisual responses could be either higher or lower than the mean auditory-alone responses, as shown by the reasonably symmetric distribution of Z-score differences in Figure 7b. There was little difference in the overall Fano factors on auditory-alone A trials vs. audiovisual B trials (Figure 7c, median auditory 1.8, median audiovisual 1.9).

**Figure 7:**
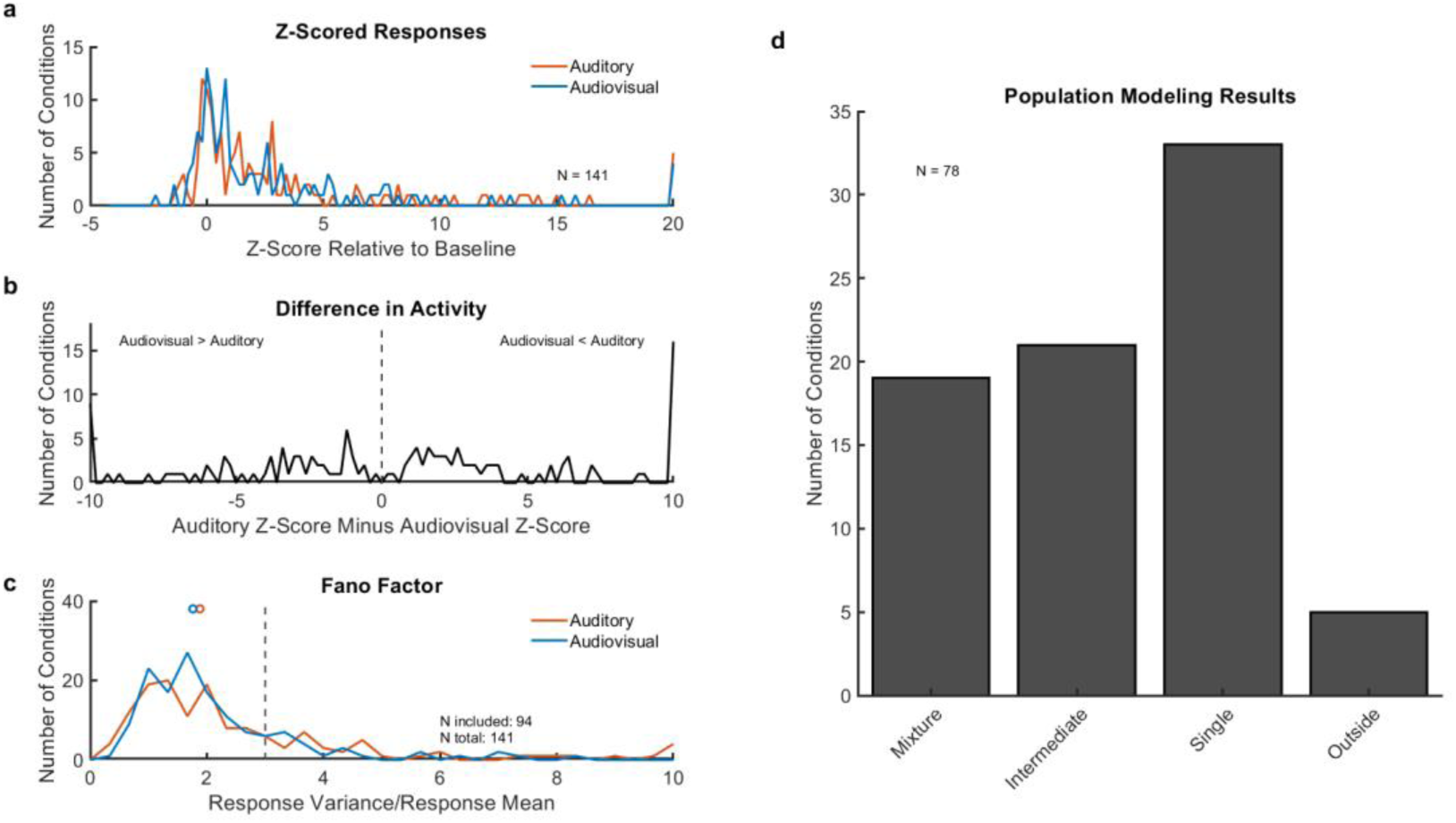
Screening and modeling results for a visual cue accompanying one of two sounds. In all panels, analysis and plotting conventions are the same as for Figure 3. a) Distribution of mean responses on the auditory-alone (red) and audiovisual-alone (blue) conditions. b) Difference between these two mean Z-scores for each triplet. N=141 showed sufficiently different levels of activity for inclusion. c) Variance-to-mean ratio (Fano factor) of the spike count responses on auditory (red) and audiovisual (blue) conditions. 94 triplets were included for further analysis. d) Results of the trial-to-trial analysis for triplets classified with a probability greater than 50%. Singles and Mixtures collectively make up 66.7% of conditions, indicating that information is preserved about a sound and a sound-light pair from trial to trial.

Across the entire population, Single was again the most common finding, occurring in 33 of 78 included triplets, or 42.3% (Figure 7d). These were evenly split between preferring the auditory stimulus (n=17) and the audiovisual one (n=16), but predominantly favored the low-frequency stimulus (n=31) over the high frequency one (n=2; see Table 1 for full breakdown). Mixtures occurred in 19 of 78, or 24.3%, about the same proportion as occurred when two sounds were presented without either being accompanied by a visual stimulus. Within the Mixture category, 2 triplets were classified as showing Slow fluctuations (Figure 8a), 7 showed Fast fluctuations (Figure 8b,c), and 7 showed Overreaching properties (Figure 8d,e), again illustrating that some of the fluctuating activity in the Mixtures was not well matched to the benchmarks (detailed categorization shown in Supplementary Figure 3). In net, though, between the continued presence of Mixtures, and the relatively balanced presence of Singles preferring either the auditory-alone or audiovisual condition, these results suggest that pairing one of two sounds with a visual stimulus doesn’t greatly change the proportion of neurons encoding that sound within the IC. We also conducted a similar analysis using two sounds as the benchmarks for responses to a sound and a sound-light pair, and results for were similar for these triplets, indicating that either a sound-light pair or its component lone sound may serve as a possible benchmark for these fluctuating responses (Supplementary Figure 4).

**Figure 8:**
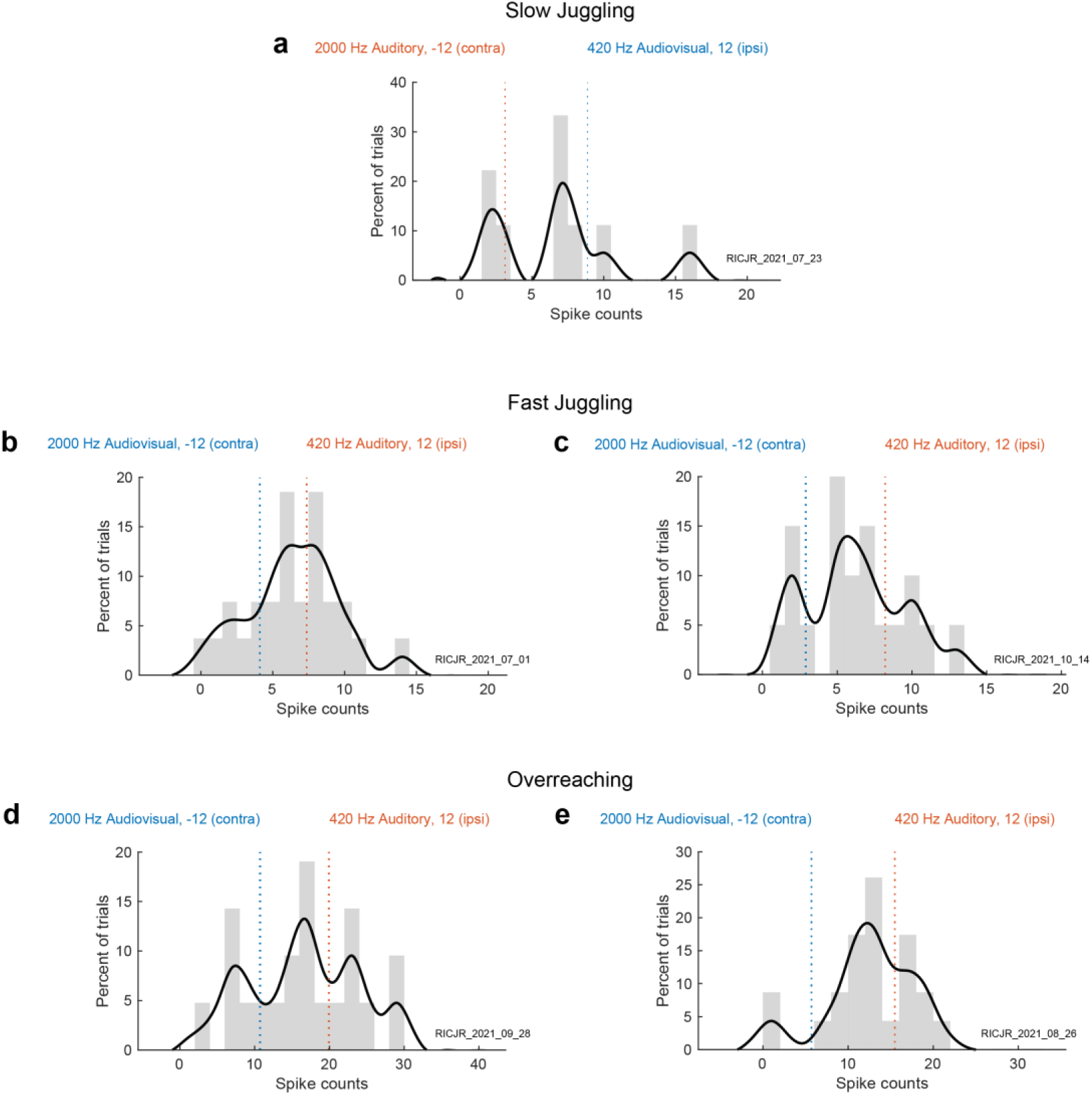
Example conditions involving a sound and sound-light pair. Example neurons’ Mixture responses (black) to individual conditions consisting of one sound (red) and one sound-light pair (blue). Analysis and plotting conventions are the same as for Figures 4-5. a) Example of Slow Juggling. b-c) Examples of Fast Juggling. d-e) Examples of Overreaching.

**Table 1:** Preferences of triplets classified as Single in the auditory-audiovisual dataset. Adding a visual stimulus did not appear to affect the preferences of individual neurons, which remained heavily biased in favor of low frequency sounds.

|  | Low<br>Frequency | High<br>Frequency |
| --- | --- | --- |
| Auditory | 15 | 2 |
| Audiovisual | 16 | 0 |

### Sound-light pairs at both locations

We also included trials in which visual stimuli were presented with both sounds. By including audiovisual conditions in both hemifields, these trials were symmetric and balanced in space, similar to the auditory-auditory trials, and serve as an interesting point of comparison to both the auditory-auditory and the auditory-audiovisual conditions (Figure 9). The population results for this dataset were overall similar to the others, except that the incidence of Mixtures was the highest – 17 of 53 or 32.1% of conditions, equal to Singles, and just ahead of Intermediates (14 of 53 or 26.4%). Most of the Mixtures were classified as Overreaching (n=12), again showing that the benchmarks do not always precisely capture the fluctuation modes, with a few classified as Fast (n=2) or Slow (n=1). Examples are shown in Figure 10.

**Figure 9:**
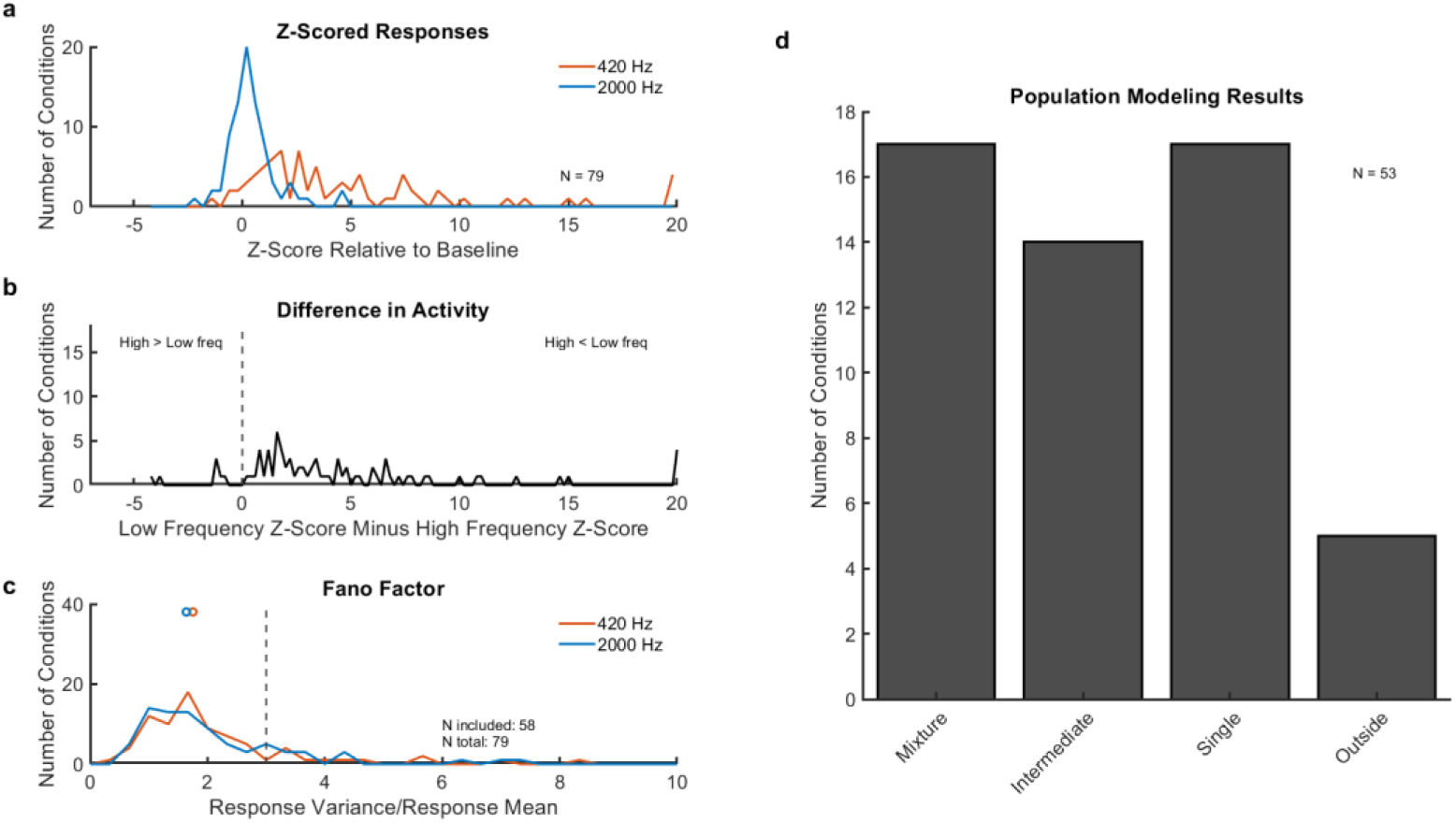
Screening and modeling results for two visual cues accompanying two sounds. In all panels, analysis and plotting conventions are the same as for Figure 3. a) Distribution of mean responses in conditions with a 420 Hz sound paired with a visual stimulus (red) and a 2000 Hz sound paired with a visual stimulus (blue). b) Difference between these two mean Z-scores for each triplet. N=79 showed sufficiently different levels of activity for inclusion. c) Variance-to-mean ratio (Fano factor) of the spike count responses in conditions with a 420 Hz sound and visual stimulus (red) and a 2000 Hz sound and visual stimulus (blue). 58 triplets were included for further analysis. d) Results of the trial-to-trial analysis for triplets classified with a probability greater than 50%. Singles and Mixtures collectively make up 64.2% of conditions, indicating that information is preserved about both sound-light pairs from trial to trial.

**Figure 10:**
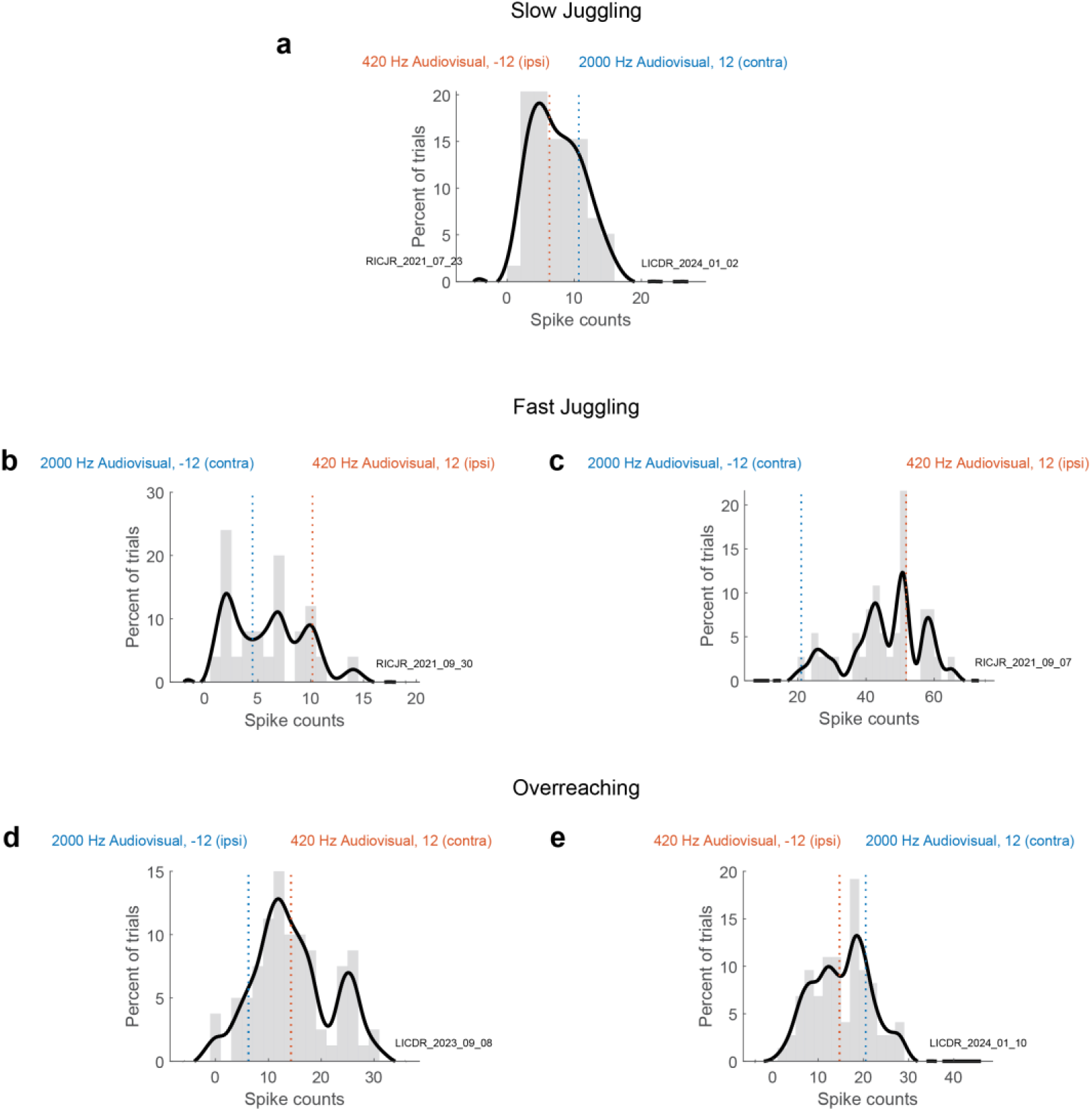
Example conditions involving two sound-light pairs. Example neurons’ Mixture responses to individual conditions consisting of a 420 Hz sound paired with a light (A) and a 2000 Hz sound paired with a light (B). Analysis and plotting conventions are the same as for Figures 4-5. a) Example of Slow Juggling. b-c) Examples of Fast Juggling. d-e) Examples of Overreaching.

### Influence of visual cues on fluctuations within individual trials

In earlier sections, we focused on neural fluctuations at relatively large time scales, spanning multiple trials or entire individual trials. Although useful in aggregate, these trial-to-trial analysis methods may neglect to uncover fluctuations at a faster time scale. Specifically, an Intermediate response might indicate averaging of the A and B responses during a trial, or it might indicate a fluctuating response at a faster time scale than the time window that we analyzed. As seen in Figures 3, 6, 7, and 9, the proportion of triplets classified as Intermediate varied across the different datasets, suggesting that indeed there could be differences in the patterns of fluctuations at the within-trial time scale. To evaluate how the neural response might evolve over the course of an individual trial, we used the Dynamic Admixture Point Process (DAPP) Model, a statistical technique previously developed and described by (Caruso et al., 2018; Glynn et al., 2021) (Model 3, Figure 1c). Briefly, the technique assesses neural responses according to their shape over the course of a trial, providing a descriptive numerical score for the degree of fluctuation, or “waviness,” over time within the trial (for details, see Materials and Methods and (Caruso et al., 2018; Glynn et al., 2021)). A higher waviness score indicates more within-trial fluctuation between the A and B responses, while a lower waviness score indicates relatively “flat” or unchanging activity.

Figure 11 shows the average numerical scores for within-trial waviness, for symmetric (auditory-auditory and audiovisual-audiovisual) and asymmetric (auditory-audiovisual) visual cue conditions. Across all conditions and neurons, fluctuations on conditions with asymmetric visual stimuli showed more waviness than fluctuations on conditions with symmetric visual stimuli. These results suggest that the presence of one lone sound and one sound-light pair induces within-trial patterns of fluctuating activity to a greater degree than is the case when there are two sounds alone or two sounds which are both paired with visual stimuli.

**Figure 11:**
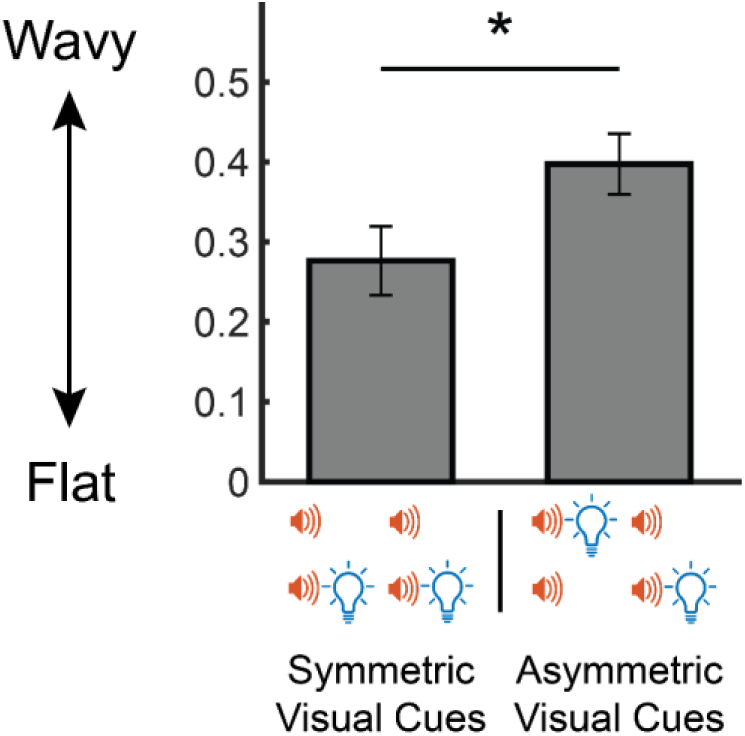
Within-trial “waviness” is higher when a visual stimulus is paired with only one of two sounds (asymmetric case) than when visual stimuli are either symmetric or absent. Scores contributing to within-trial waviness across all triplets screened for a winning probability greater than 0.5 and A and B Fano factors less than 3, showing the comparison between conditions with no visual stimuli or visual stimuli on both sides (symmetric) and conditions involving visual stimuli paired with just one sound (asymmetric). Conditions with asymmetric visual stimuli tend to be wavier, indicating a response that fluctuates more during the course of individual trials (two-sample t-test, p = 0.0380).

Together, these results demonstrate that there are differences in within-trial activity in the IC depending on the number of visual stimuli that accompany two simultaneous sounds.

### Relationship between behavioral and neural biases

The findings reported above suggest that when two sounds are present, both sounds are represented in the IC even when one or more sounds are paired with a visual stimulus (although this representation may involve shifted benchmarks in some cases). But they do not preclude the possibility that there might be subtle biases in the representation which might correlate with biases in the performance of the behavioral task. For example, visual cues might create a bias at the level of individual trials, causing neurons that fluctuate to spend more time (that is to say, trials) responding to one sound than the other. Such shifts in preference are not discernable from the classification tasks carried out above. For example, the Mixture category could include a symmetric 50-50 split of A-like and B-like trials, but it could also include an asymmetric 80-20 split between the two types. Therefore, further analysis is needed to discern if pairing sounds with light might cause such shifts in preference between the stimuli. Any such underlying bias toward one of the component stimuli might in turn be related to the animal’s perception and/or behavior. In the case of trials with one sound and one sound-light pair, more trials might respond to the visually-paired sound than to the unpaired sound, or trials might favor the sound that the monkey localizes first on each trial.

To address these possibilities, we developed a method to describe which stimulus is favored during each single trial, using a trial-wise score indicating which of the two targets is best represented in the neural response (Figure 12a). This analysis was run on auditory-audiovisual conditions that were classified as Mixtures. For each auditory-audiovisual trial, a Bayes classifier was used to determine how well the spike count on that trial matched either the corresponding single-location auditory-alone or the single-location audiovisual-alone response distributions, yielding a probability score between 0 (audiovisual) and 1 (auditory-only). A similar process can be used with the one-location labels assigned differently, such as according to their location (−12 or +12 degrees), sound frequency (420 or 2000 Hz), or saccade order (which target was localized first vs. second).

**Figure 12:**
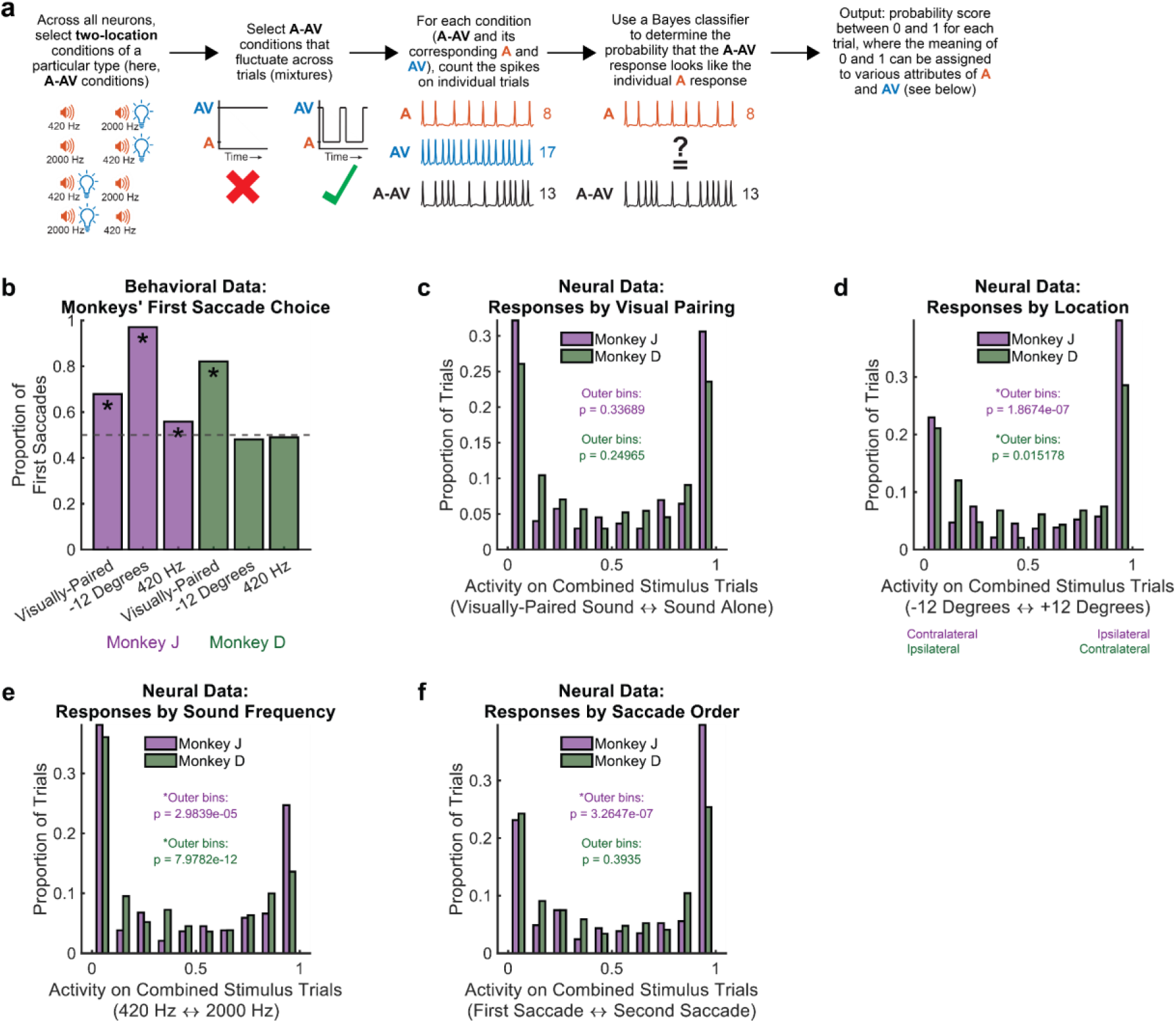
When two sounds are present and one sound is paired with a visual stimulus, neural activity has an ambiguous relationship with behavior. a) Schematic representing the analysis process for a score that indicates which target (lone sound, A, or sound-light pair, AV) is best represented in the neural data on an individual trial. A Bayes classifier is used to compare the spike counts on two-location trials (A-AV) to the spike counts on trials that present each target individually (A or AV alone). The output from the analysis is a probability score between 0 and 1 for each two-location trial, indicating which single target is best represented in the neural response. b) Behavioral tendencies of Monkey J (purple bars) and Monkey D (green bars) to direct their first saccade to targets defined by three stimulus attributes: visual pairing, location, and sound frequency. Asterisks indicate proportions that differ from 0.5 (binomial test, p<0.05). c) Distribution of trials best responding to each target on two-location trials, defined according to which sound is visually-paired. Trial-wise scores on the x-axis are calculated as described in (a). Each data point is an individual trial for conditions that had asymmetric visual cues, showed a fluctuating response from trial to trial (Mixture), and had A and B Fano factors less than 3. 1016 trials across these selected conditions met the inclusion criteria (Monkey J: 575 trials, purple bars; Monkey D: 441 trials, green bars). Neurons in both monkeys are equally likely to respond to the visually-paired vs. lone sound on individual trials (binomial test on outer bins; Monkey J: purple bars, p > 0.05; Monkey D: green bars, p > 0.05). d) Same as (c) but with the trial-wise score defined according to target location. Neurons in both monkeys spend more trials responding to the sound located at 12 degrees (binomial test on outer bins; Monkey J: purple bars, p < 0.05; Monkey D: green bars, p < 0.05). e) Same as (c) but with the trial-wise score defined according to sound frequency. Neurons in both monkeys spend more trials responding to the 420 Hz sound (binomial test on outer bins; Monkey J: purple bars, p < 0.05; Monkey D: green bars, p < 0.05). f) Same as (c) but with the trial-wise score defined according to which target the monkey directed the first vs. second saccade during the trial. Neurons in Monkey J spend more trials responding to the target that is localized second, while neurons in Monkey D, represent both saccade targets equally (binomial test on outer bins; Monkey J: purple bars, p < 0.05; Monkey D: green bars, p > 0.05).

To interpret these scores, we first considered each monkey’s general behavioral patterns. While the behavioral task itself involved no particular incentive structure in terms of which location the monkey might saccade to first, the monkeys could in principle exhibit consistent performance biases. For example, they might tend to look first to the visually-paired sound, to the stimulus on the left or right, or to the lower vs. higher frequency sound. These behavioral tendencies might correlate with each monkey’s neural activity on individual trials. Figure 12b shows the tendency of each monkey to direct the first saccade on each trial toward one component vs. the other on these two-location trials, with consideration for the visual pairing, location, and frequency of each sound. Asterisks represent proportions that differ from 0.5 (binomial test, p<0.05). Different behavioral strategies emerged for each monkey: while Monkey J typically localized the target at −12 degrees first (contralateral to the recorded IC for that monkey), Monkey D was more likely to localize the visually-paired sound first, regardless of its location or frequency.

Figures 12c-f explore the distributions of neural activity scores across all trials, with scores assigned according to different attributes of the two target stimuli. Note that in all of these cases, peaks are expected near both 0 and 1 due to the selection for Mixtures. We are therefore interested in whether one peak is larger than the other, whether this trend is specific to each monkey, and whether the neural activity relates to that monkey’s behavioral biases.

In Figure 12c, the scores correspond to which target was visually-paired during each trial. Neither monkey shows a neural bias toward one target over the other (binomial test on outer bins; Monkey J: purple bars, p > 0.05; Monkey D: green bars, p > 0.05). This is true despite Monkey D having a behavioral preference to localize the visually-paired sound first. In Figure 12d, the scores correspond to the location of each target. Both monkeys display a neural bias toward the target located at 12 degrees (binomial test on outer bins; Monkey J: purple bars, p < 0.05; Monkey D: green bars, p < 0.05), although this bias opposes Monkey J’s behavioral preference and is unrelated to Monkey D’s behavioral preference. In Figure 12e, the scores correspond to the frequency of each sound. Both monkeys display a neural bias toward the 420 Hz target (binomial test on outer bins; Monkey J: purple bars, p < 0.05; Monkey D: green bars, p < 0.05), consistent with the low-frequency bias identified in Figure 6. However, only Monkey J showed any difference in behavioral performance as a function of sound frequency, and this effect was modest (Figure 12b).

The preceding analysis considered the relationship between neural and behavioral biases on average. What about trial-wise correlation? Given that monkeys showed variation in which stimulus they localized first, it’s possible that the responses on those trials could be correlated with the behavioral sequence on that specific trial. Supplementary Figure 5 shows an additional way of visualizing these results on a trial-wise basis, both pooled across monkeys (a-c), and separated by monkey (d-i). In each panel, each dot represents the activity of a neuron on a particular trial, plotted as a function of where the monkey directed the first saccade.

Supplementary Figure 5a shows this comparison based on whether the monkey’s first saccade was to the visually-paired sound (left column) or to the unpaired sound (right column). The neural activity scores are the same as in Figure 12c, representing which sound was favored in the neural response on each trial. In this panel, on average, the neural activity appeared more similar to the sound-alone activity on trials when the monkeys looked at the visually-paired sound first – as if the IC was holding a representation of the *second* saccade target location. This difference is statistically significant (Mann-Whitney U-test, p < 0.05). This pattern held across visual pairing, location (Supplementary Figure 5b, with neural activity scores corresponding to Figure 12d), and frequency (Supplementary Figure 5c, with neural activity scores corresponding to Figure 12e), but was only significant when the data were broken down based on visual pairing and frequency. The pattern is largely driven by the results of Monkey J (d-f), and was not statistically significant for Monkey D (g-i).

Figure 12f quantifies this pattern in histogram form similar to Figure 12c-e, showing the scores corresponding to which target the monkey directed the first vs. second saccade during each trial. This panel serves as a summary that combines the data from the previous two figures, providing a trial-wise comparison between a monkey’s behavior and the corresponding neural activity during each trial. While Monkey D shows no neural bias, Monkey J displays a strong neural bias to respond to the second localized target (binomial test on outer bins; Monkey J: purple bars, p < 0.05; Monkey D: green bars, p > 0.05).

In summary, neural activity in this task appears to have a complex relationship to a monkey’s behavior. Monkey J employs a consistent behavioral strategy of localizing the leftward target first (Figure 12b), yet the neural activity typically favors the rightward (ipsilateral) target (Figure 12d), which is localized second (Figure 12f). Similarly, Monkey D employs a behavioral strategy of localizing the visually-paired sound first (Figure 12b), yet the neural activity displays no such trend (Figure 12c). Instead, Monkey D’s neural responses favor the rightward (contralateral) target (Figure 12d) or the 420 Hz sound (Figure 12e), regardless of which of these is localized first (Figure 12f).

## Discussion

How the brain represents more than one sensory signal at a time is uncertain. One well-understood mechanism is for different sensory signals to be processed by different populations of neurons. This can occur in a variety of ways. In the visual system, specialization in the form of receptive fields or tuning for particular stimulus characteristics leads to different populations responsive to different stimuli. However, such selectivity is not thought to be sufficient to fully “sort” different stimuli onto distinct and separate populations of neurons. In most visual areas, receptive fields are broader than would seem necessary to support the spatial acuity that is evident in perceptual testing (for review, see (Groh et al., 2024)). And in most (perhaps all) mammalian auditory brain regions, circumscribed auditory receptive fields do not exist at all (Groh et al., 2003; McAlpine & Grothe, 2003; Porter & Groh, 2006; Woods et al., 2006; Werner-Reiss & Groh, 2008; Salminen et al., 2009, 2010; Grothe et al., 2010). Rather, sound location appears to be rate coded, involving varying firing rates in a fairly uniform population of neurons (Middlebrooks et al., 1994, 1998; Groh et al., 2003; Zwiers et al., 2004; Werner-Reiss & Groh, 2008; Salminen et al., 2009; Lee & Groh, 2014). These issues have led to the proposal of time-domain multiplexing as an additional means of ensuring the representation of multiple sensory signals despite overlap in the population of neurons potentially responsive to each sensory input (Caruso et al., 2018; Mohl et al., 2020; Glynn et al., 2021; N. Y. Jun et al., 2022; Schmehl et al., 2024; Chen et al., 2024; Groh et al., 2024).

In this context, what happens when there are signals from two different sensory modalities is particularly interesting. At the input stages, different sensory signals are separate from each other – photoreceptors in the eyes and hair cells in the ears process visual and auditory input separately. But later in the brain, the sensory systems can potentially interact with one another (Bulkin & Groh, 2006; Ghazanfar & Schroeder, 2006; Kayser & Logothetis, 2007; Gruters & Groh, 2012; Teichert & Bolz, 2018; Schmehl & Groh, 2021). Here, we investigated these interactions in a predominantly auditory structure, the inferior colliculus, and evaluated what happens when visual stimuli are paired with sounds.

Our first finding is that even when there is only one visual stimulus and only one sound, and they are presented at the same location in space, fluctuating activity is elicited in a subpopulation of neurons. This finding is important for two reasons. First, it indicates that even though IC neurons can be affected by visual input (Porter et al., 2007; Bulkin & Groh, 2012; Schmehl et al., 2025), there appears to be a mechanism by which the visual and auditory inputs are kept more or less distinct. This is different from the common view of what is referred to as “multisensory integration,” in which responses to multimodal inputs are seen as being combined to yield novel firing rates (Meredith & Stein, 1986; Wallace et al., 1998; Stanford et al., 2005; Alvarado et al., 2007). Second, it indicates that fluctuating activity can occur even when only one stimulus *location* is involved. Previous studies identifying such multiplexing patterns have all involved stimuli at separate locations, raising the possibility that spatial attention might be fluctuating between the two locations. That fluctuating activity can be observed even when two stimuli are at the same location illustrates that spatial attention cannot be the only explanation for such activity patterns.

Our second main question was whether and how visual stimuli might influence previously observed patterns of fluctuation when two sounds are present (Caruso et al., 2018). Here, we mainly find that the kinds of fluctuations elicited by two sounds continue to occur when one or both sounds are paired with a visual stimulus. This differs from related work in auditory cortex, which suggested that pairing a visual stimulus with one of two sounds can cause auditory responses to be “captured” by the paired visual stimulus (Atilgan et al., 2018). While these differences may relate to species or methodological differences (the auditory cortical work is in ferrets and involved sounds varying in frequency over time), they do suggest that inferior colliculus and auditory cortex may play different roles. The IC appears capable of preserving information about all sounds, regardless of the presence of visual stimuli. Such a capacity is likely needed in the brain to ensure that auditory perception can unfold without interference from visual inputs, which may not always be beneficial. For example, the visual scene at any given moment may consist of objects irrelevant to auditory processing, and some method of ensuring that sound processing is unaffected by visual signals is therefore needed.

When comparing neural responses with behavior, the patterns are nuanced. On the whole, both stimuli are represented, even though monkeys can localize the stimuli in either order. When evaluating correlations at the trial-wise level, one monkey shows no correlation and the other shows a pattern in which the stimulus that the monkey localized second is slightly more represented than the stimulus it localized first. Together, these patterns are consistent with a view of the IC as representing both stimuli, regardless of the behavioral pattern. These findings are in contrast with a previous study that used a similar task structure (Caruso et al., 2018), which concluded that the initially localized target was slightly overrepresented in the neural activity. Several factors may account for these contrasting findings. As shown in Figure 12, monkeys may adopt unique behavioral preferences during the task. The monkeys in this experiment were different from those in (Caruso et al., 2018), suggesting that a variety of behavioral strategies and neural relationships may be possible across subjects. These differences may underlie differences in sound localization ability or sensory binding across individuals, and further studies involving more subjects could shed additional light on these possibilities.

On the whole, the weak relationship between behavioral and neural biases is actually consistent with the overall conceptual framing of the study: some areas of the brain should process all stimuli, with selection for those of particular behavioral relevance occurring later (Broadbent, 1957). If the IC is one of the “process it all” brain regions, then any connection to behavioral performance is likely to be weak and variable. As the IC is fundamentally an auditory structure, perhaps adding a visual cue to a sound is sufficient to modulate a neuron’s firing rate ((Schmehl et al., 2025) and Figure 3b), but not sufficient to influence how a monkey distinguishes among more than one sound to support behavior.

Limitations and directions for future inquiry include the various screening steps that restrict which triplets can be tested. Some of these screenings are inevitable. For example, the requirement that the responses to the two stimuli differ from one another is necessary from a theoretical perspective: if the single stimulus spike count benchmarks are indistinguishable from one another, then it is impossible to determine whether combinations of stimuli evoke fluctuations between those benchmarks with any confidence (see (Mohl et al., 2020) for a sensitivity analysis). In short, some screening is unavoidable, and multiplexing analysis will always be restricted to investigating this property in a subset of neurons.

The screening on the basis of the Poisson assumption is a more interesting case. The models used in the current study are formulated in relation to the expected variability of Poisson distributions. A screen on the single stimulus response distributions is needed to ensure that this assumption is not dramatically violated when only one stimulus is presented. Future work will be needed to consider alternative distributions that may sometimes provide a better fit to the single stimulus response distributions, such as a negative binomial distribution. A statistical approach that does not employ a Poisson assumption is currently under development (Marco et al., 2025). The Marco et al. method also introduces analysis at the level of spike trains/interspike intervals, thus providing finer temporal resolution and overcoming the limitations inherent in the spike-count based methods used here.

Another consideration involves the benchmarking on the basis of single-stimulus response patterns. As illustrated by the results using the SCAMPI modeling approach, not all fluctuating activity patterns are well captured as alternation between firing patterns well matched to the A and B response distributions. One possible explanation for this may be that the A and B response distributions themselves include fluctuating activity patterns – perhaps due to encoding signals other than the sensory stimuli deliberately presented in the experimental paradigm (for discussion, see (Groh et al., 2024; Chen et al., 2024)). This is especially likely to be a factor for the analysis of the auditory-audiovisual and audiovisual-audiovisual triplets – since the audiovisual combinations themselves show fluctuating activity, this can be present within in these larger combinations of stimuli. At this time, we lack statistical methods for testing this hypothesis; future work will be needed.

More broadly, these findings suggest that fluctuating activity in the IC is amenable to multisensory influences. In combination with prior studies of fluctuating activity involving different types of objects (Schmehl et al., 2024), these results contribute to a growing body of evidence suggesting that fluctuating activity may be a general strategy for preserving information about more than one object, even if one of those objects is not preferred by the neuron under study (Li et al., 2016; Caruso et al., 2018; Kienitz et al., 2018, 2022; N. Y. Jun et al., 2022; Schmehl et al., 2024; Groh et al., 2024). Additionally, intriguing questions remain concerning how these responses may be coordinated across multiple neurons. We have previously identified both positive and negative response correlations in a similar multiplexing analysis involving simultaneously recorded V1 neurons (N. Y. Jun et al., 2022), suggesting a form of coordination that ensures that both stimuli are represented across the population. Whether and how such multi-neuron coordination occurs in the IC is not yet known, and progress toward studying this possibility has historically been limited by the technical challenges associated with conducting multi-unit recordings in deep brain structures. New technologies and tools are rapidly developing and could facilitate experiments that probe this question in detail (J. J. Jun et al., 2017; Hall et al., 2021). Future work that explores the simultaneous responses of multiple IC neurons could clarify how neurons coordinate their responses to sounds in the presence of visual input.

How a multiplexed code may be created by upstream circuitry and read-out by downstream circuitry remains unknown as well, but some theoretical possibilities have been proposed (Caruso et al., 2018; Groh et al., 2024). The critical elements will undoubtedly involve selective switching such that not all inputs/outputs to a given neuron are “open” for through traffic at all times – inputs and outputs must be gated, and they may or may not be controlled by some top-down supervisory clock cycle to coordinate this switching process.

Together, these results suggest that visual stimuli can influence the IC’s responses to complex soundscapes. This neural finding may underlie behavioral phenomena such as visual capture, in which a salient visual stimulus can alter perception of a sound’s attributes, particularly the sound’s location (Thomas, 1941; Jackson, 1953; Thurlow & Jack, 1973; Bertelson & Radeau, 1981; Alais & Burr, 2004; Atilgan et al., 2018). A greater understanding of how visual stimuli inform the representation of multiple sounds in the IC, a relatively early sensory area, helps illuminate how the brain uses multisensory information to aid the perception of complex scenes across the sensory hierarchy.

## Acknowledgements

The authors would like to thank members of the Groh Lab and Multiplexing Working Group, including Nicholas Marco, Shawn Willett, Jeff Mohl, Valeria Caruso, Justine Shih, Tingan Zhu, Chadbourne Smith, Stephanie Lovich, and Cynthia King, for their feedback on this work. This work was supported by the National Institutes of Health (F31DC020361 to MNS, R01NS129112 to JMG and STT, R01DC016363 to JMG and STT, and R01DC013906 to JMG and STT).

## Supplementary Figures

**Supplementary Figure 1:**
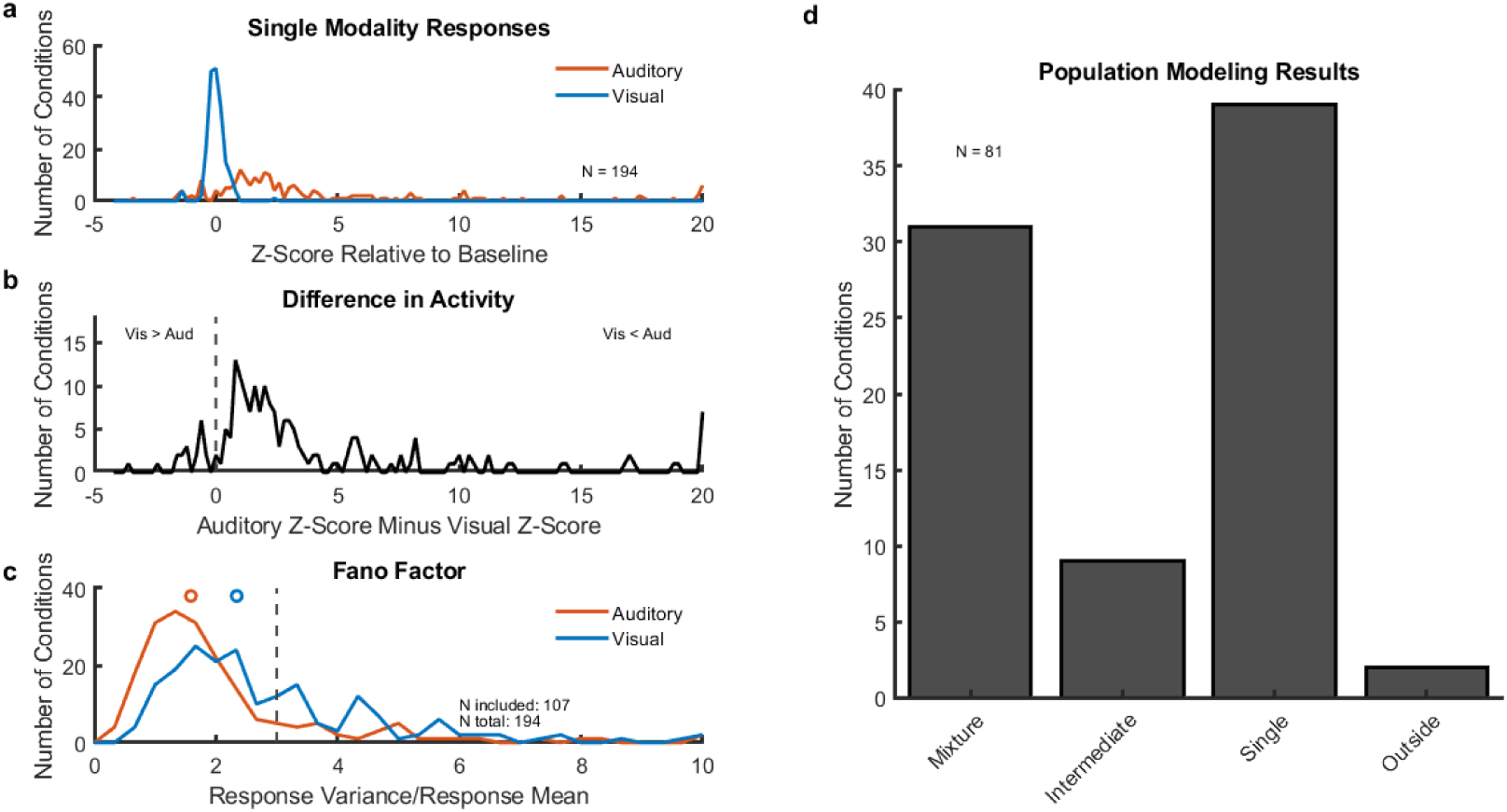
Screening and modeling results for sound-light pairs analyzed 100-500 ms following stimulus onset show that the proportion of mixtures is comparable when the spike counting window is delayed past the onset latency of the visual responses. Same as Figure 3 but responses are analyzed 100-500 ms following stimulus onset, thus beginning after the longest visual response latency in this dataset. In all panels, analysis and plotting conventions are otherwise the same as for Figure 3. a) Distribution of mean responses on the auditory (red) and visual (blue) conditions. b) Difference between these two mean Z-scores for each triplet. A total of 194 triplets showed sufficiently different levels of activity for inclusion. c) Variance-to-mean ratio (Fano factor) of the spike count responses on auditory (red) and visual (blue) conditions. After Fano factor screening (Fano < 3, dashed line), 107 triplets were included for further analysis. d) Results of the trial-to-trial analysis for triplets classified with a probability greater than 50% (N=81 triplets). Mixtures make up 31 of 81 total, or 38.3%, higher than the 30 of 92 (32.6%) for the 0-500 ms spike counting window in Figure 3. Singles and Mixtures collectively make up 86.4% of conditions, indicating a high degree of trial-to-trial fluctuation between the single auditory and single visual responses.

**Supplementary Figure 2:**
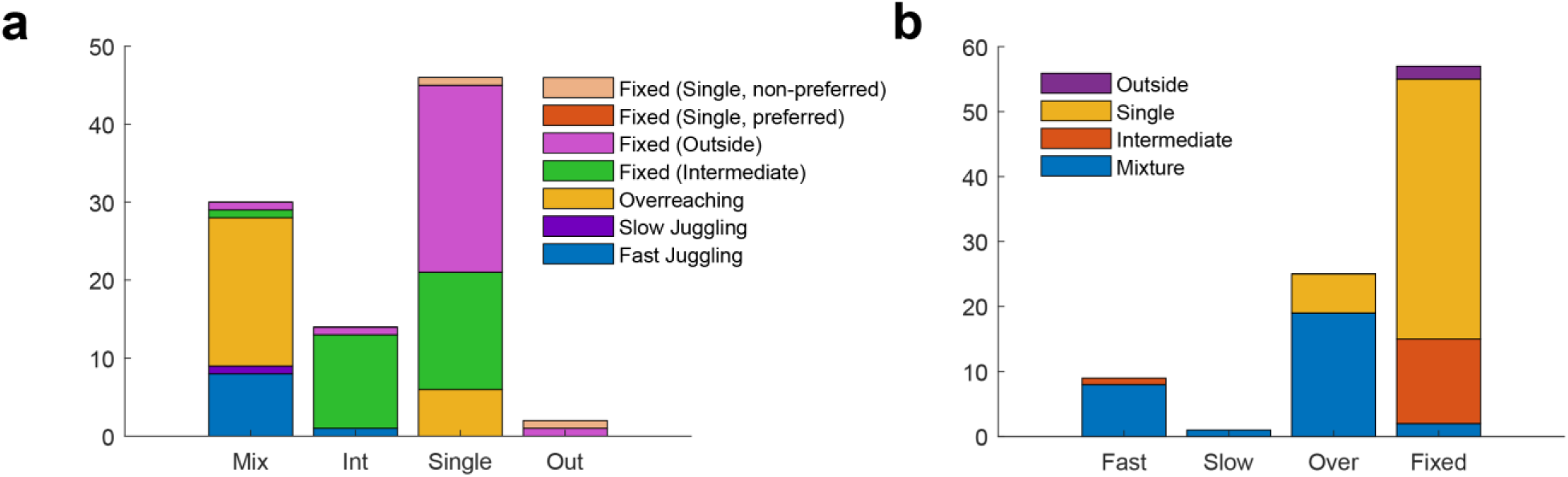
Full correspondence of results of Model 1 and Model 2 (SCAMPI) for a sound-light pair. a) Model 1 results split into Model 2 results. b) Model 2 results split into Model 1 results. Generally, the assignments in both panels correspond in the expected fashion, such as most Mixtures are assigned to Fast Juggling, Slow Juggling, or Overreaching as described in the main text, and Intermediates are generally categorized as Fixed (Intermediate). Model 2 re-categorized some Singles as Fixed (Outside), suggesting that the original Single category was too generous and missed some cases in which the visual stimulus caused an increase in activity when combined with a sound. Such cases are described more fully in (Schmehl et al., 2025).

**Supplementary Figure 3:**
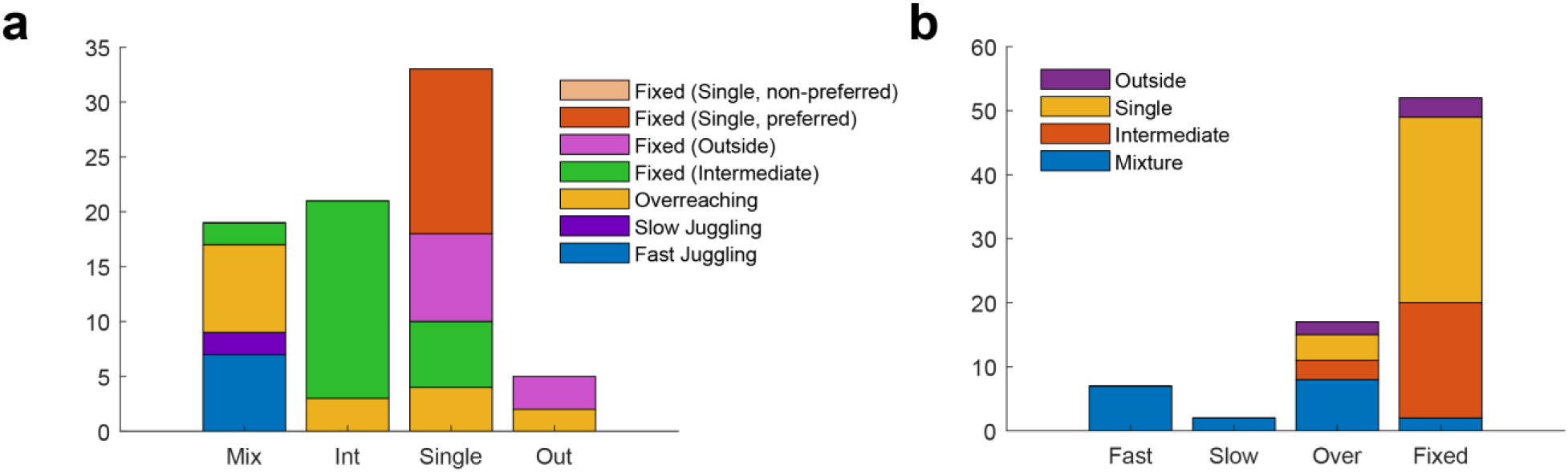
Full correspondence of results of Model 1 and Model 2 (SCAMPI) for auditory-audiovisual conditions. a) Model 1 results split into Model 2 results. b) Model 2 results split into Model 1 results. As in Supplementary Figure 2, the assignments in both panels are generally as expected, with the addition of some Overreaching in the Intermediates.

**Supplementary Figure 4:**
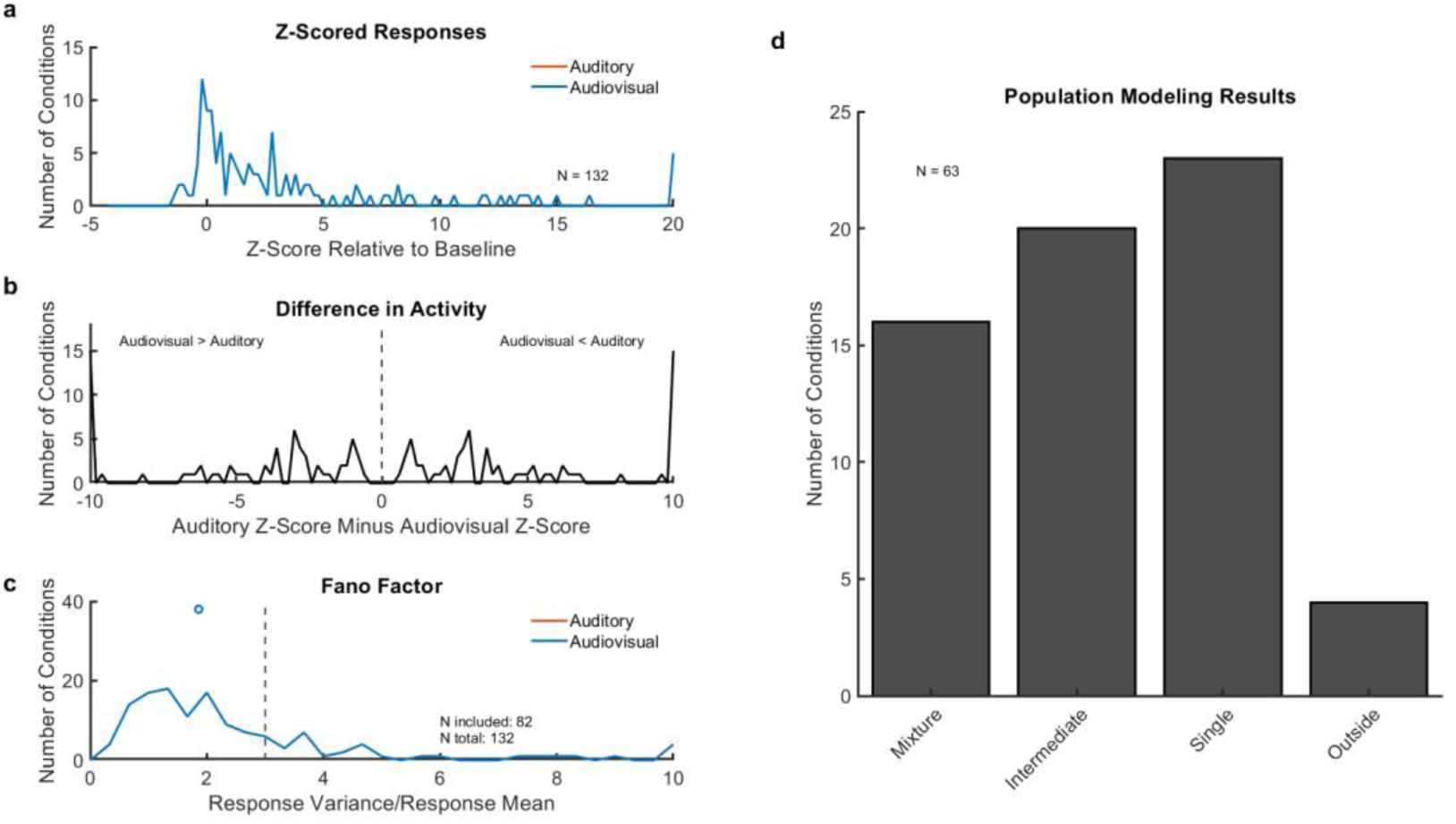
Screening and modeling results when two lone sounds are used as benchmarks for a visual cue accompanying one of two sounds. Same as Figure 7 but when responses are benchmarked to a sound (A) and a sound that is presented with a visual stimulus in the AB condition (B) – i.e., “audiovisual” indicates the sound that is visually-paired in AB, but here we are benchmarking to the responses to that sound alone. In all panels, analysis and plotting conventions are the same as for Figure 3. a) Distribution of mean responses on the auditory-alone and audiovisual sound presented alone trials (both blue; note that A and B in these triplets are counterbalanced and so the distributions are identical and overlapping). b) Difference between these two mean Z-scores for each triplet. N=132 showed sufficiently different levels of activity for inclusion. c) Variance-to-mean ratio (Fano factor) of the spike count responses on auditory (red) and audiovisual sound alone (blue) trials. 82 triplets were included for further analysis. d) Results of the trial-to-trial analysis for triplets classified with a probability greater than 50%. Singles and Mixtures collectively make up 61.9% of conditions, indicating that information is preserved about a sound and a sound-light pair from trial to trial, even when responses are benchmarked to each sound alone.

**Supplementary Figure 5:**
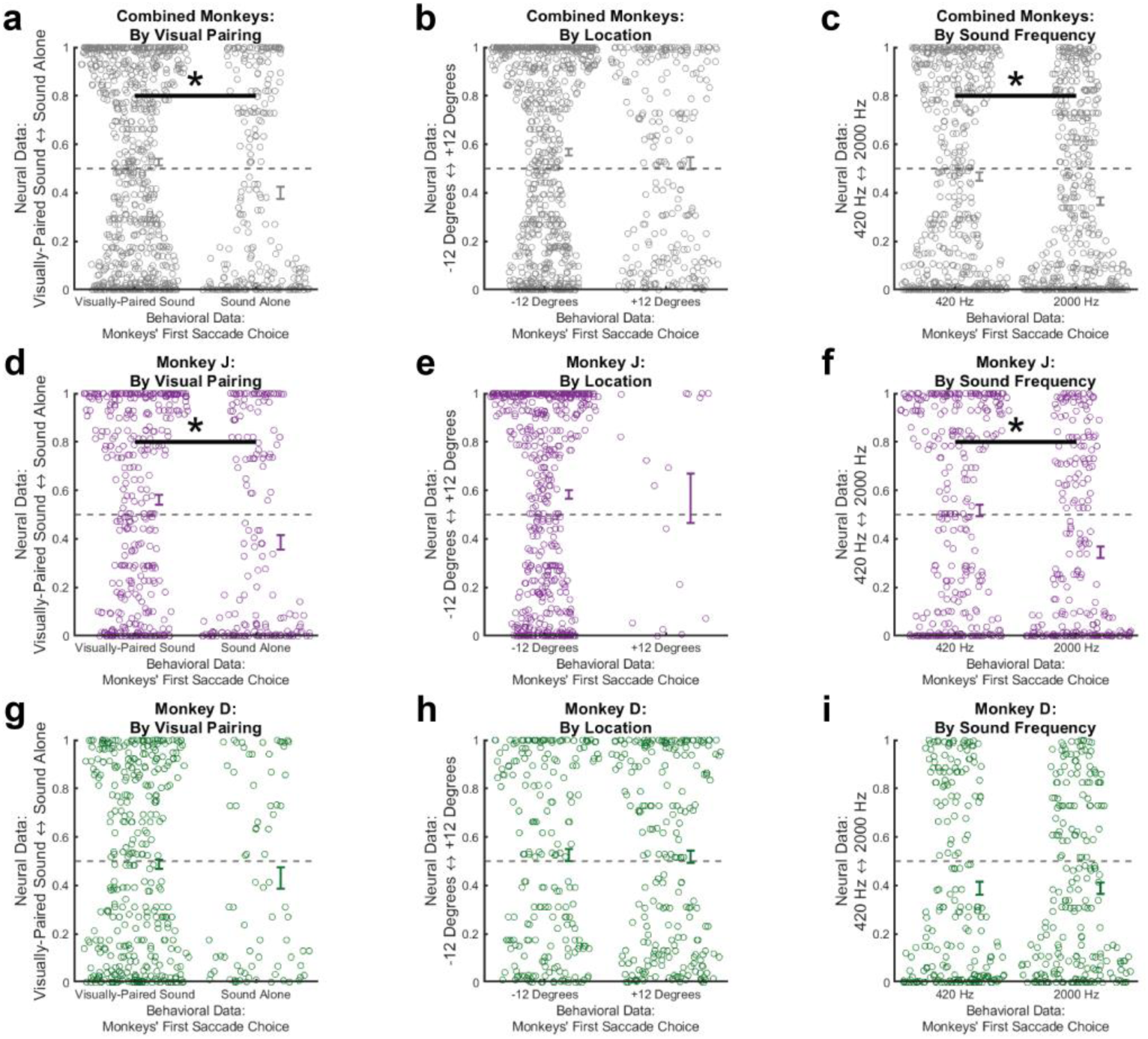
Trial-wise neural activity and relationship with behavior. a) Comparison of neural activity scores on individual trials, split by trials when the monkey made the first saccade to the visually-paired sound or the sound alone. Neural activity is more likely to correlate with the target that was localized second (Mann-Whitney U-test, p < 0.05). b) Same as (a) but with the trial-wise score defined according to target location. Neural activity is roughly equally biased toward each sound regardless of the monkeys’ behavioral choice (Mann-Whitney U-test, p > 0.05). c) Same as (a) but with the trial-wise score defined according to sound frequency. Neural activity is more likely to correlate with the target that was localized second (Mann-Whitney U-test, p < 0.05). d-f) Same as (a-c) but for Monkey J only. The same trend holds where a significant difference in neural activity emerges when the data are organized by visual pairing and frequency. g-i) Same as (a-c) but for Monkey D only. No significant relationship is evident between neural activity and behavior.

